# PRMT1/Hmt1 drives α-synuclein aggregate dissolution through a catalysis-independent pathway

**DOI:** 10.64898/2026.02.27.708458

**Authors:** Sakshi Dewasthale, Purusharth I Rajyaguru

## Abstract

Protein homeostasis, the balance between protein folding, function, and clearance, is essential for cellular health, and its disruption is a hallmark of many neurodegenerative diseases, including Parkinson’s disease. Arginine methylation, carried out by enzymes like Hmt1 in yeast and PRMT1 in humans, is best known for regulating transcription, translation, apoptosis and pre-mRNA splicing. In this study, we reveal a surprising, non-canonical role for these methyltransferases in controlling the aggregation and toxicity of α-synuclein, a protein lacking arginine and implicated in Parkinson’s disease pathology. We find that under oxidative stress, Hmt1 and PRMT1 relocate from the nucleus into the cytoplasm. Using genetic, biochemical and imaging assays in yeast and mammalian cells, we observe that Hmt1/PRMT1 dissolve aggregates, promotes their removal through the ubiquitin–proteasome system, and reduces α-synuclein toxicity. Importantly, this activity does not require the classical methyltransferase function. Catalytically inactive mutants and a specific PRMT1 inhibitor both enhance aggregate clearance and improve cell survival. Together, our findings uncover an unexpected function for Hmt1 and PRMT1 in maintaining protein quality under stress, expanding our understanding of how cells cope with toxic protein assemblies and pointing to new directions for therapeutic strategies in protein aggregation disorders.

## Introduction

Protein activity and abundance in the cell are fine-tuned through posttranslational modifications (PTMs) ^1^. One such PTM, arginine methylation, is evolutionarily conserved yet comparatively less understood despite its growing importance in the regulation of gene expression ^2^. The reaction is catalysed by protein arginine methyltransferases (PRMTs), which transfer methyl groups from the donor molecule S-adenosyl methionine (SAM) to guanidino nitrogen atoms of arginine residues, producing S-adenosylhomocysteine (SAH) as a byproduct^3^. Depending on the site and number of methyl additions, four distinct modifications are generated: monomethyl arginine on the terminal ω-nitrogen (ω-MMA), asymmetric dimethyl arginine (aDMA; two methyl groups on the same ω-nitrogen), symmetric dimethyl arginine (sDMA; methyl groups on each ω-nitrogens), and δ-monomethyl arginine (δ-MMA; methylation at the δ-nitrogen) ^3^.

In budding yeast, *Saccharomyces cerevisiae*, the primary enzyme catalysing this reaction is Hmt1 (HnRNP MethylTransferase) ^4^. Its homolog in higher eukaryotes, PRMT1, represents the major type I PRMT in humans and is strongly associated with the modification of RNA-binding proteins (RBPs) ^5^. Indeed, RBPs are the largest group of proteins known to undergo arginine methylation ^6^. Within these proteins, arginine–glycine–glycine (RGG) motifs serve as the preferred methylation hotspots ^7^. RGG motifs are critical elements that facilitate both protein–protein and protein–RNA interactions ^7^. While Hmt1 itself resides predominantly in the nucleus^8^, its substrates are distributed in both nucleus and cytoplasm ^9^. Importantly, many RBPs targeted by Hmt1 are enriched in cytoplasmic RNA granules such as P-bodies and stress granules, where they exert key roles in controlling mRNA metabolism and turnover ^10, 11^.

The regulation of Hmt1 partitioning between nucleus and cytoplasm remains poorly understood. We hypothesised that localization of Hmt1 would change under specific conditions and this would alter the availability of this enzyme towards its substrates which in turn could modulate the arginine methylation proteome. As Hmt1 also interacts with proteins which are not arginine methylated ^12^, we hypothesised that Hmt1 could possibly also have catalytic activity-independent functions. A notable genetic link has been identified between Hmt1 and α-Synuclein: while α-Synuclein overexpression in yeast promotes cytotoxicity, this effect is markedly intensified in Δhmt1 ^13^. In this report we identify a non-canonical role of Hmt1/ PRMT1 in disassembling α-synuclein aggregates, suggestive of a therapeutic potential of PRMT1 in pathogenesis of Parkinson’s disease.

## Results

### Hmt1 delocalises from nucleus to cytoplasm under oxidative stress

We tested localization of Hmt1 under different stress conditions, such as-heat shock (42°C for 30 mins), genotoxic stress (0.2M hydroxyurea for 90 mins), nitrogen deprivation for 30 mins, amino acid deprivation for 30 mins, oxidative stress (4mM H_2_O_2_ for 30 mins and 0.5% sodium azide for 30 mins), and glucose deprivation for 30 mins (Figure 1A). Strikingly, Hmt1 undergoes a marked loss of nuclear localization and relocalizes to cytoplasm in response to two distinct oxidative stresses such as 0.5% sodium azide for 30 mins (Figure 1A & C) and 4mM H_2_O_2_ for 30 mins (Figure 1A&D). Cytoplasmic localization is also accompanied by assembly into granules. Nuclear delocalization is reversible as the localization of Hmt1 to nucleus is restored upon 30 mins of recovery (Figure 1A). Hmt1 levels decreased upon treatment with sodium azide but increased upon H_2_O_2_ stress (Figure 1B). Strikingly, Hmt1 levels increased upon heat stress, HU, nitrogen starvation and amino acid starvation, but these stresses did not lead to nuclear delocalization and cytoplasmic localization of Hmt1 suggesting that increased protein levels likely do not contribute to this phenotype (Figure 1A &B). Endogenously tagged Hmt1 GFP also delocalised from the nucleus and localized to cytoplasm and granules upon treatment with 0.5% sodium azide for 30 mins (Supplementary Figure 1A&B) and 4mM H_2_O_2_ for 30 mins (Supplementary Figure 1A&C), suggesting that the observed phenotype is not specific to plasmid borne Hmt1.

**Figure 1:**
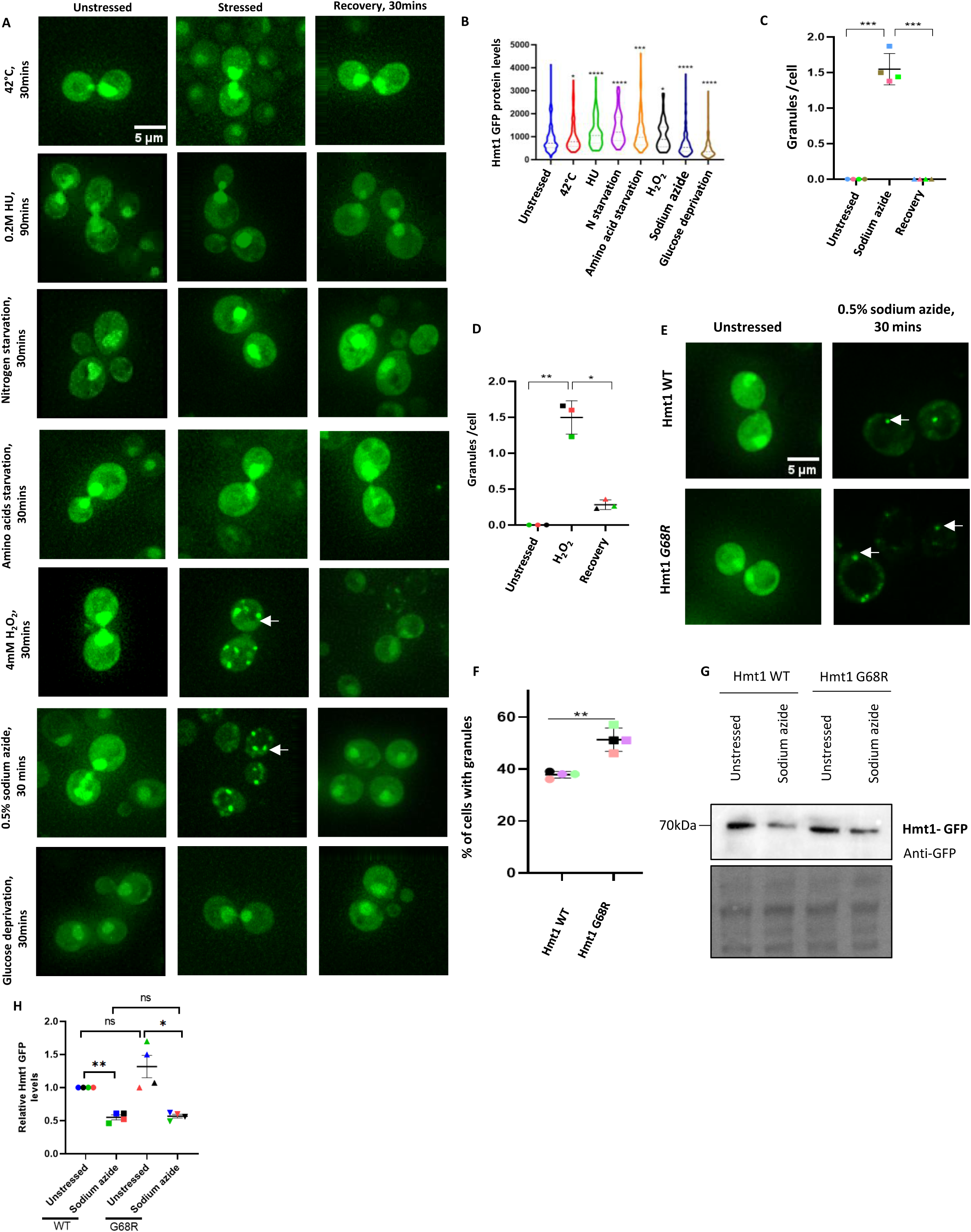
Hmt1 delocalises from nucleus to cytoplasm under oxidative stress. **A**. Live cell imaging showing localization of Hmt1 upon various stress. **B**. Quantitation for total protein levels for data presented in A. **C**. Quantitation for data presented in A upon sodium azide. **D**. Quantitation for data presented in A upon H_2_O_2_. **E.** Live cell microscopy for Hmt1 WT and G68R under unstressed and sodium azide treated conditions. **F**. Quantitation for data presented in E as % of cells with granules. **G.** Western blotting analysis to check Hmt1 GFP protein levels for Hmt1 WT and G68R under unstressed and sodium azide stressed conditions. **H**. Quantitation for data represented in G. The significance of the data was calculated by paired t-test, and P-values are summarized as follows: ***P < 0.005; **P < 0.01; *P < 0.05.

To further understand the impact of catalytic activity on localization, we tested the catalytically dead mutant of Hmt1-G68R, which is defective in binding to S-adenosyl methionine (SAM). Western blotting using anti-mono methylarginine specific antibody confirmed the lysate from Hmt1 G68R mutant did not show detectable cross-reactivity to mono-methyl arginine antibody (Supplementary Figure 1D).

The catalytically inactive Hmt1 G68R mutant also relocalized from the nucleus to the cytoplasm and, notably, showed a modest but significant increase in granule formation compared to wild-type Hmt1 (Figure 1E&F). Total protein analysis revealed comparable levels of Hmt1 WT and G68R under both unstressed and sodium azide–treated conditions, although protein levels of both forms decreased upon stress (Figure 1G&H). These observations indicate that stress-induced cytoplasmic relocalization of Hmt1 occurs independently of its catalytic activity.

We further observed that there was no colocalization between Hmt1 granules and the granules of its cytoplasmic substrates, Sbp1, Scd6 and P-body marker Edc3 (Supplementary Figure 1E), indicating that the cytoplasmic granules of Hmt1 are likely not sites of arginine methylation.

The above observations suggest that Hmt1 localization to granules is specific to oxidative stress and a potentially non-canonical function. Oxidative stress in cells is known to cause protein misfolding and damage^14^. We therefore think that Hmt1 could be acting as a part of protein quality control pathway, as a part of its non-canonical function.

### Hmt1 regulates α-synuclein toxicity

Yeast has been used as a model organism to study the physiology of α-synuclein. Interestingly, a genetic interaction between Hmt1 and α-Synuclein has been reported. Deletion of HMT1 enhances the toxicity of α-Synuclein ^13^. There are no arginine residues in α-synuclein. We therefore hypothesised that Hmt1 could be involved in regulating the toxicity of α-synuclein in catalytic activity independent manner.

To address this hypothesis, we first checked the toxicity of α-Synuclein in Δhmt1 cells which enhanced the synuclein toxicity as compared to wild type. Δhmt1 does not have an inherent growth defect as EV transformed strains of WT and Δhmt1 cells grew comparable to each other (Figure 2A). To test if enhanced toxicity was dependent on the catalytic activity we complemented Δhmt1 strain with Hmt1 wild type and Hmt1-G68R (catalytically dead mutant defective in binding SAM). We observed that the G68R mutant rescued the toxicity better than Hmt1 WT (Figure 2B). Analysis of the protein levels in these strains indicated that the toxicity was correlated to α-synuclein protein levels. α-synuclein levels were highest in Δhmt1 which showed the highest growth defect. Complementation of Δhmt1 with Hmt1 WT, reduced α-synuclein to levels comparable to WT whereas complementation of Δhmt1 with Hmt1 G68R reduced the protein levels more as compared to wild type Hmt1 (Figure 2C&D). These results suggest that Hmt1 is to be involved in maintaining the protein levels of α-synuclein independent of its catalytic activity.

**Figure 2:**
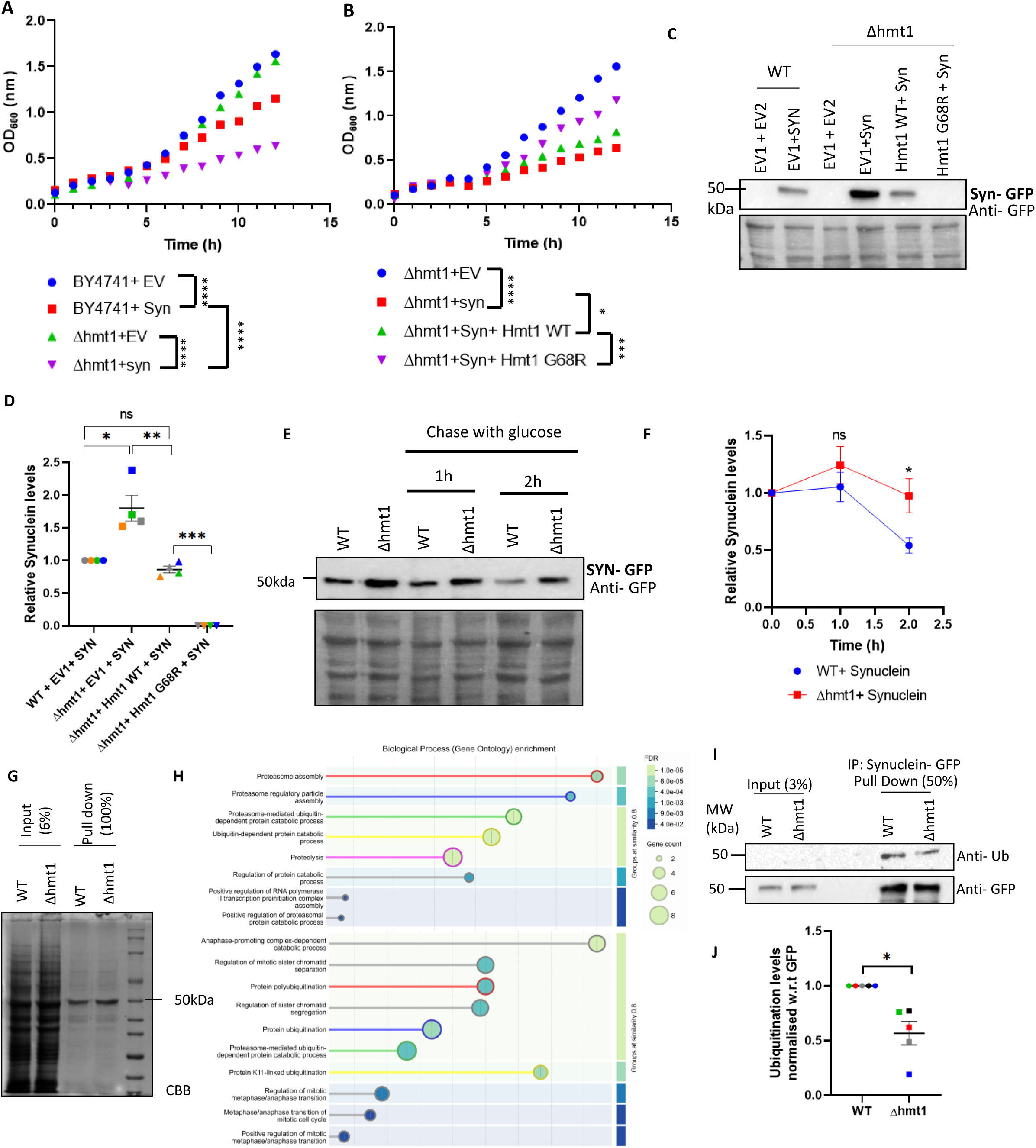
Hmt1 regulates synuclein toxicity and protein levels. **A**. Growth curve analysis upon synuclein overexpression in WT and Δhmt1 strains. Significance for growth curve in A, calculated by two-way ANOVA. **B.** Growth curve analysis with synuclein overexpression upon complementing Δhmt1 with Hmt1 WT and G68R mutant. Significance for growth curve in B, calculated by two-way ANOVA. **C.** Western blotting analysis to check the levels of synuclein in different strains. **D**. Quantitation for synuclein-GFP levels. Ponceau serves as loading control. **E**. Western blotting analysis to check SYN protein levels after shifting cultures to glucose containing media, showing slower degradation in the absence of Hmt1. Ponceau serves as loading control. **F**. Quantitation for E, by two-way ANOVA. **G.** Pull down for SYN from WT and Δhmt1 background. **H**. Gene ontology (GO) analysis from mass spectroscopy data for proteins interacting with SYN specifically in WT and absent in Δhmt1, showing enrichment of proteins associated with the Ubiquitin-proteasome machinery. **I.** Pull down for Synuclein GFP to check ubiquitination status. **J**. Quantitation for data in I. The significance of the data was calculated by paired t-test, and P-values are summarized as follows: ***P < 0.005; **P < 0.01; *P < 0.05.

Interestingly deletion of two other known arginine methyltransferases-Sfm1 and Rmt2, did not affect α-synuclein toxicity compared to WT, indicating that this effect was specific to Hmt1 (Supplementary Figure 3A).

### α-synuclein protein degradation is defective in the absence of Hmt1

To understand the role of Hmt1 in regulating α-synuclein protein levels, we measured the levels of galactose-induced α-synuclein protein levels by shifting cultures to glucose containing media (Figure 2E). Degradation of α-synuclein was slower in *Δhmt1* cells as compared to WT (Figure 2E&F), indicating a role of Hmt1 in mediating α-synuclein degradation.

α-synuclein levels are known to be regulated by autophagy and ubiquitination ^15^. To understand the role of Hmt1 in autophagy mediated α-synuclein degradation, we treated cells with 100nM Rapamycin, a potent inducer of autophagy, for 90 minutes. DMSO served as vehicle control. Upon normalizing the α-synuclein levels in rapamycin with respect to DMSO, we did not observe any change in α-synuclein protein levels between WT and Δhmt1, suggesting that Hmt1 is likely not involved in autophagy-mediated α-synuclein degradation (Supplementary Figure 4).

To test the involvement of proteasomal machinery in mediating α-synuclein degradation, we performed α-synuclein pull down followed by mass spectrometry from WT and Δhmt1 cells (Figure 2G). We observed proteins associated with ubiquitin-proteasome system, specifically interacting with α-synuclein in WT and not in Δhmt1 (Figure 2H). This suggests that Hmt1 might be involved in ubiquitin mediated α-synuclein degradation.

To test this hypothesis further, we checked ubiquitination status by pulling down α-synuclein from WT and Δhmt1 cells (Figure 2I). We observed that α-synuclein was significantly less ubiquitinated in Δhmt1 as compared to WT (Figure 2I&J), suggesting that Hmt1 is likely involved in regulating α-synuclein levels by ubiquitin-mediated degradation. No ubiquitination signal was observed in Δubi4, which codes for ubiquitin, suggesting that the signal was specific to ubiquitination (Supplementary Figure 2E&F).

### α-synuclein granules increase in Δhmt1 cells

As increased protein levels of α-Synuclein were observed in Δhmt1 cells, we wanted to check if Hmt1 could also regulate the localization of α-Synuclein to granules which could depend on protein levels. Therefore, we checked the granule localization of α-synuclein under oxidative stress-4mM H_2_O_2_ for 30 mins. We observed increased α-synuclein granule assembly in Δhmt1 strain compared to WT under unstressed conditions (Figure 3A&B). Treatment with H_2_O_2_ lead to an increase in α-synuclein granules for both WT and Δhmt1 with latter forming more α-synuclein granules compared to WT upon H_2_O_2_ treatment (Figure 3A&B). Interestingly Δhmt1 also showed more α-synuclein granules than WT under recovery conditions, indicating Hmt1 might be involved in the disassembly of α-synuclein granules (Figure 3A&C). Δhmt1 also showed increase in α-synuclein protein levels under unstressed, stressed and recovery conditions compared to WT cells (Figure 3D&E).

**Figure 3:**
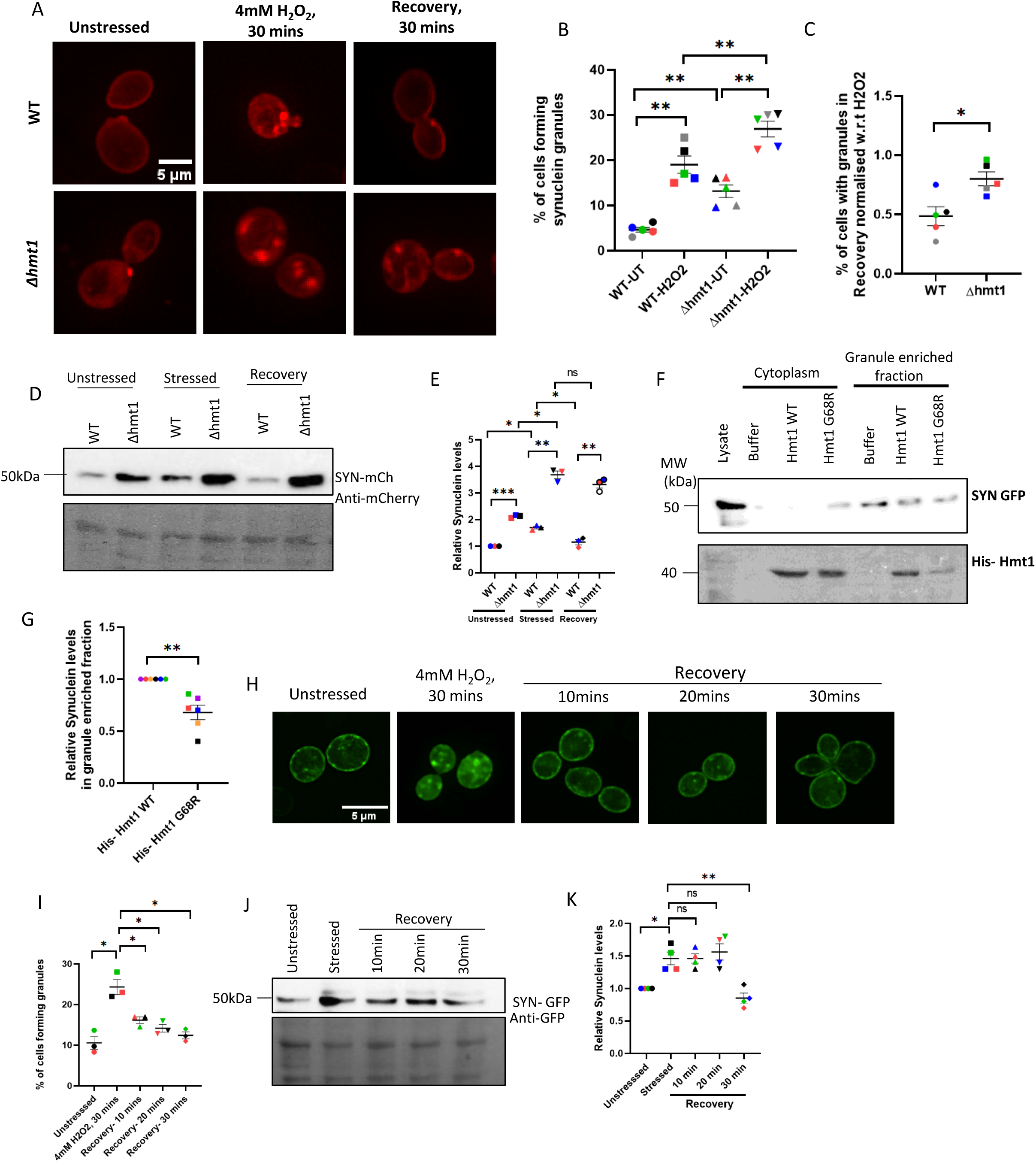
Hmt1 acts as a disassembly factor for Synuclein granules. **A.** Live cell microscopy for synuclein-mCherry in WT and Δhmt1 strains under unstressed, H_2_O_2_ and recovery conditions. **B**. Quantitation for A in terms of % of cells forming granules. **C.** Quantitation for A as % of cells forming granules upon recovery normalized w.r.t H_2_O_2_. **D**. Western blotting analysis to check Synuclein levels in WT and Δhmt1 strains under unstress, H_2_O_2_ stress and recovery conditions. **E.** Quantitation for data represented in D. **F.** Granule enrichment for synuclein upon addition of recombinant-purified Hmt1. **G**. Quantitation for synuclein present in granule enriched fraction. **H.** Microscopy for Synuclein under unstressed, stressed and recovery conditions. **I.** Quantitation for H in terms of % of cells forming granules. **J**. western blotting analysis to check total protein levels of SYN under unstressed, stressed and recovery conditions. **K**. Quantitation for data represented in J. The significance of the data was calculated by paired t-test, and P-values are summarized as follows: ***P < 0.005; **P < 0.01; *P < 0.05.

Deletion of two other known arginine methyltransferases-Sfm1 and Rmt2, did not affect α-synuclein granules under unstress and stress conditions compared to WT, indicating that this effect was specific to Hmt1 (Supplementary Figure 3B&C).

### Hmt1 acts as a disassembly factor for α-synuclein granules

To understand if Hmt1 could directly disassemble α-synuclein granules, we performed granule enrichment for α-synuclein. Compared to buffer control, addition of recombinant-purified Hmt1 WT reduced α-synuclein in the granule-enriched fraction, indicating that Hmt1 can disassemble α-synuclein granules. This reduction was further enhanced with the Hmt1 G68R mutant (Figure 3F&G).

Hmt1 was responsible for α-synuclein degradation as well as granule disassembly, we hypothesised that disassembly could precede degradation. To address this, we monitored recovery at multiple time points following 30 min H₂O₂ stress. α-synuclein granules began to disassemble within 10 min of recovery (Figure 3H&I), whereas total α-synuclein protein levels decreased only after 30 min (Figure 3J&K). These results indicate that granule disassembly precedes the reduction in α-synuclein protein levels.

### PRMT1 suppresses α-synuclein granule formation in HeLa cells

To check if the disassembly of α-synuclein granules by Hmt1 is conserved, we extended our findings to HeLa cells. We performed immunocyto-chemistry for PRMT1 to check the localization of PRMT1 upon oxidative stress. We observed that PRMT1 (like Hmt1) localised mostly to the nucleus under unstressed conditions and de-localised from the nucleus and formed granules in the cytoplasm upon treatment with 0.5mM sodium arsenite and 2mM H_2_O_2_ for 1h (Figure 4A, B&C). The total protein levels of PRMT1 decreased upon treatment with sodium arsenite and H_2_O_2_, suggesting that localization to granules is not due to increased protein levels (Figure 4D).

**Figure 4:**
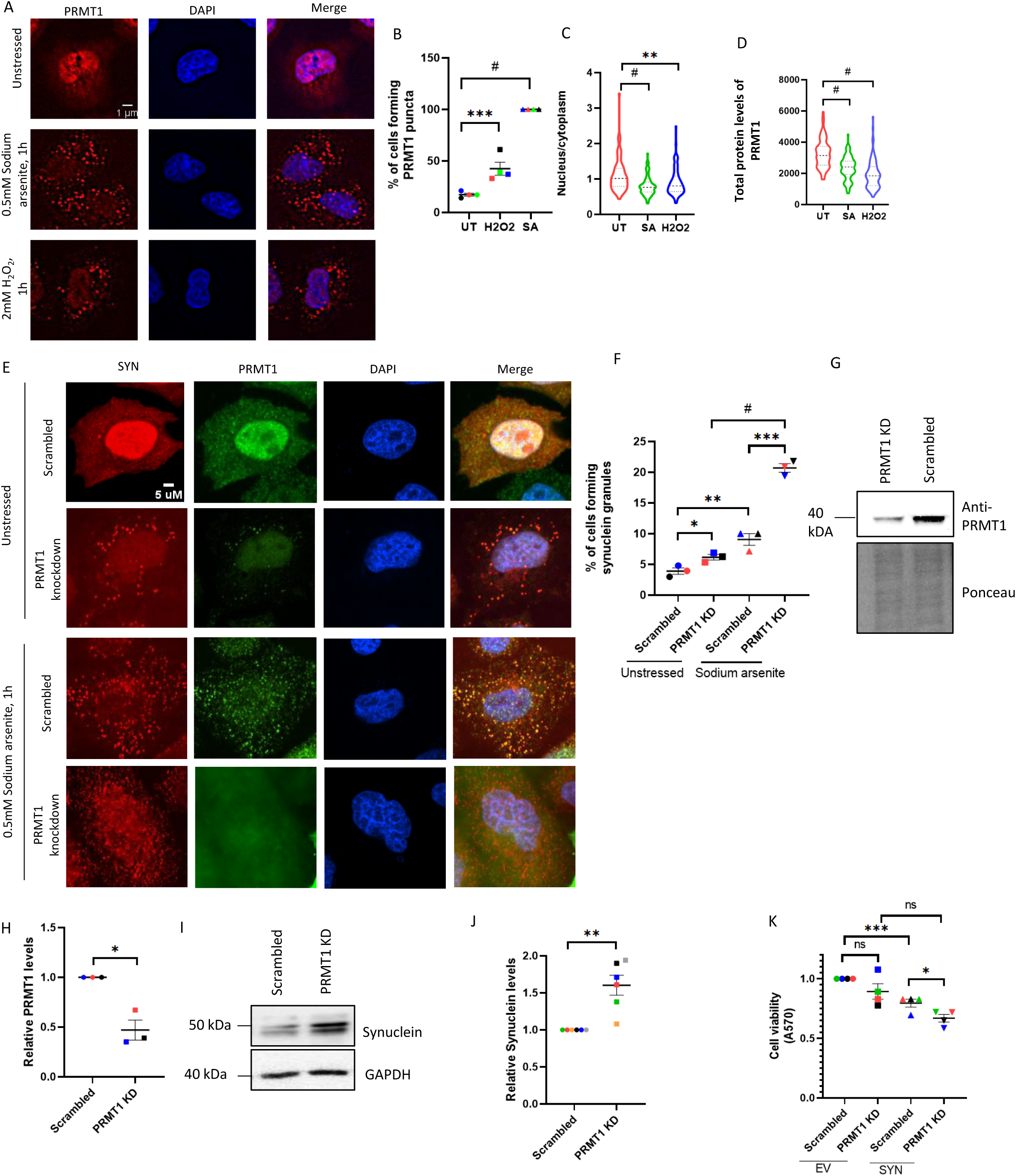
PRMT1 suppresses α-synuclein puncta formation in HeLa cells. **A.** ICC using Anti-PRMT1 antibody to check localization of PRMT1 under unstressed, sodium arsenite and H_2_O_2_ treated conditions. **B**. Quantitation for A in terms of % of cells forming granules **C.** N/C ratio, **D.** total protein levels of PRMT1. **E.** Microscopy for SYN upon PRMT1 knockdown. **F.** Quantitation for data represented in E as % of cells forming SYN granules. **G**. Western analysis for confirming PRMT1 knockdown. **H**. Quantitation for data represented in G. **I**. Western analysis for checking Synuclein levels upon PRMT1 knockdown. **J**. Quantitation for data represented in I. **K**. MTT assay for checking SYN toxicity upon PRMT1 knockdown. The significance of the data was calculated by paired t-test, and P-values are summarized as follows: ***P < 0.005; **P < 0.01; *P < 0.05.

To assess whether PRMT1 affects α-synuclein localization to granules, we overexpressed PRMT1 and observed a reduced percentage of cells containing α-synuclein granules under both unstressed conditions and following 0.5 mM sodium arsenite treatment (Supplementary Figure 5A&B). Consistently, western blot analysis showed that PRMT1 overexpression also reduced total α-synuclein protein levels (Supplementary Figure 5C&D).

To further examine the role of PRMT1 in regulating α-synuclein aggregation, we performed shRNA-mediated knockdown of PRMT1. PRMT1 depletion increased α-synuclein granules (Figure 4E&F) and total α-synuclein protein levels (Figure 4I&J) under both unstressed and sodium arsenite–treated condition. PRMT1 knockdown was validated by western blotting, showing ∼50% reduction in PRMT1 protein (Figure 4G&H). Consistent with observations in the Δhmt1 strain, PRMT1 knockdown enhanced α-synuclein–associated toxicity in HeLa cells (Figure 4K), while PRMT1 knockdown alone did not affect cell viability (Supplementary Figure 6A).

### PRMT1 acts as a disassembly factor for α-synuclein aggregates in HeLa cells

To test whether PRMT1 can disassemble α-synuclein granules, recombinant-purified PRMT1–GST was incubated with α-synuclein lysate, followed by granule partitioning. Compared to buffer control, PRMT1 addition reduced α-synuclein in the granule-partitioned fraction with a concomitant increase in the cytoplasmic fraction, indicating granule disassembly. GAPDH served as a negative control and showed no enrichment in the granule-partitioned fraction (Figure 5A&B).

**Figure 5:**
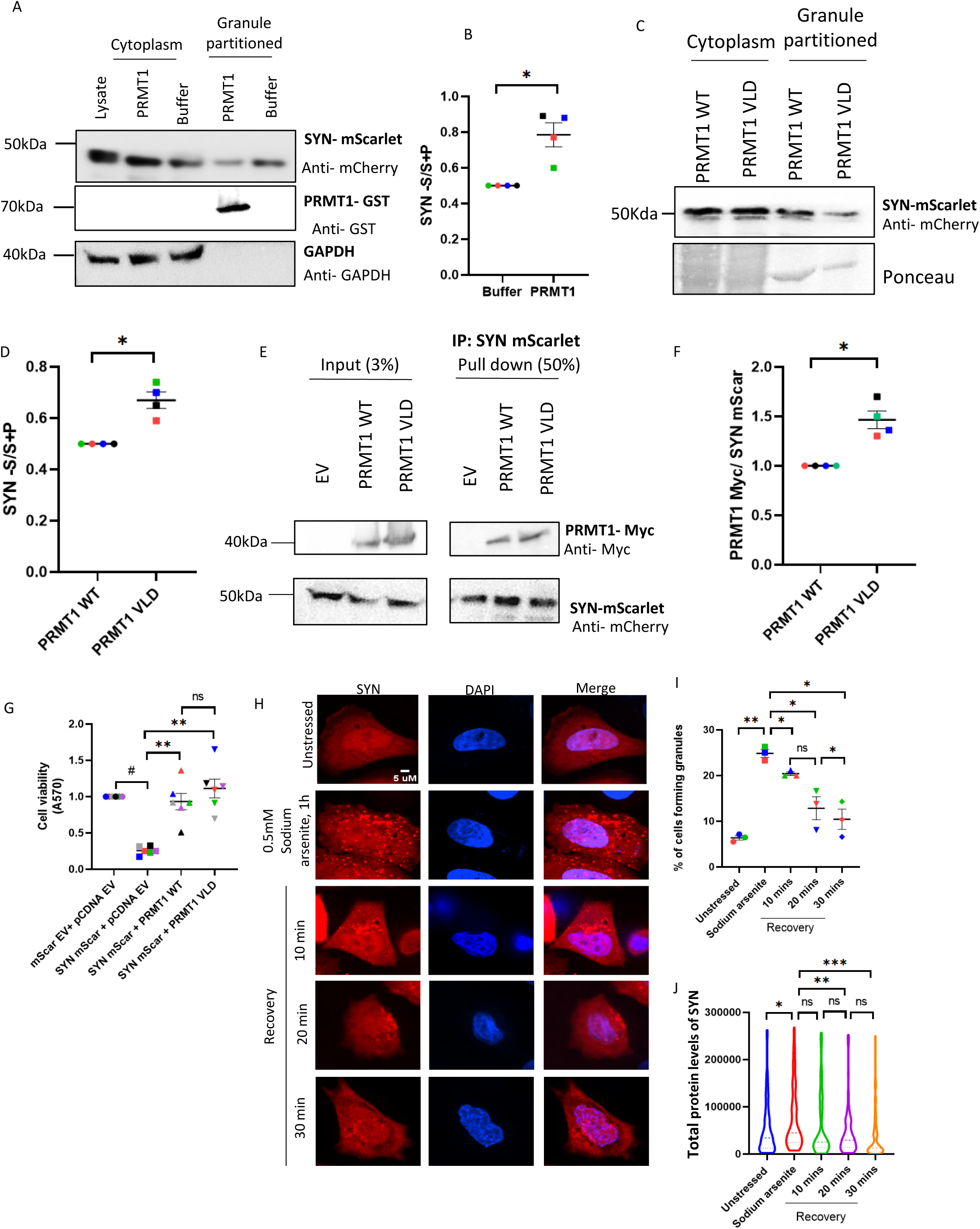
PRMT1 acts as a disassembly factor for Synuclein in HeLa cells. **A.** Granule enrichment for synuclein upon addition of recombinant-purified PRMT1. **B.** Quantitation for data represented in A. **C.** Granule enrichment for synuclein upon addition of PRMT1 WT and PRMT1 VLD expressing cell lysates. **D.** Quantitation for data represented in C. **E**. SYN pull down for checking SYN-PRMT1 interaction. **F.** Quantitation for E. **G.**. MTT assay for Synuclein in the presence of PRMT1 WT and catalytically dead mutant. **H**. Microscopy for Synuclein under unstressed, stressed and recovery conditions. **I.** Quantitation for data represented in H as% of cells forming granules. **J**. Analysis of total protein levels of SYN under unstressed, stressed and recovery conditions. **I**. The significance of the data was calculated by paired t-test, and P-values are summarized as follows: ***P < 0.005; **P < 0.01; *P < 0.05.

To compare the disassembly activities of PRMT1 WT and the catalytically inactive, SAM-binding mutant-PRMT1 VLD, α-synuclein lysate were incubated with lysates expressing PRMT1 WT or PRMT1 VLD, followed by granule partitioning. PRMT1 VLD reduced α-synuclein partitioning into granules more effectively than PRMT1 WT, indicating enhanced granule disassembly by the mutant (Figure 5C&D).

To test if the disassembly of synuclein granules was mediated via interaction with PRMT1 we performed α-synuclein pull down from HeLa cells and observed that α-synuclein interacted with PRMT1 WT, whereas PRMT1 VLD mutant interacted relatively better with α-synuclein (Figure 5E&F).

Overexpression of α-synuclein leads to aggregation which is toxic^16^. Therefore it would be expected that disassembly of α-synuclein aggregates could alleviate toxicity. Consistently, MTT assays showed reduced viability of HeLa cells upon α-synuclein overexpression (Figure 5G). Co-expression of PRMT1 rescued this toxicity, and notably, the catalytically dead PRMT1 mutant provided a comparable rescue, indicating a catalytic activity–independent function of PRMT1 (Figure 5G). Expression of PRMT1 WT or mutant alone did not affect cell viability relative to empty vector controls, confirming that the observed effects were specific to α-synuclein (Supplementary Figure 6B).

### α-synuclein granule disassembly precedes decrease in protein levels in HeLa cells

PRMT1 was responsible for α-synuclein degradation as well as granule disassembly, we hypothesised that disassembly could precede degradation. Cells were allowed to recover for indicated times following treatment with 0.5mM sodium arsenite for 1h. α-synuclein granules began to disassemble within 10 min of recovery (Figure 5H&I), whereas total α-synuclein protein levels decreased only after 20 min (Figure 5H&J), indicating that disassembly precedes degradation.

### PRMT1 inhibitor (TCE5003) decreases α-synuclein aggregates in HeLa cells

Since the catalytically dead PRMT1 mutant showed enhanced α-synuclein binding and disassembly compared to PRMT1 WT, we hypothesized that pharmacological inhibition of PRMT1 catalytic activity would similarly reduce α-synuclein granules and rescue associated toxicity. We used the PRMT1 inhibitor TC-E-5003 which binds to the SAM binding pocket and specifically inhibits the catalytic activity of PRMT1 without affecting PRMT1 protein levels ^17^. Addition of 5µM TC-E-5003 for 3h reduced α-synuclein granules in HeLa cells under sodium arsenite treated conditions (Figure 6A&B). Western blotting analysis revealed that TC-E-5003 did not affect the total protein levels of PRMT1 (Figure 6C&D). Strikingly, addition of TC-E-5003 rescued the α-synuclein-associated toxicity (Figure 6E). Overall, our results suggest that Hmt1 and its human ortholog, PRMT1, can disassemble α-synuclein granules and further reduce α-synuclein protein levels, leading to decreased toxicity (Figure 7).

**Figure 6:**
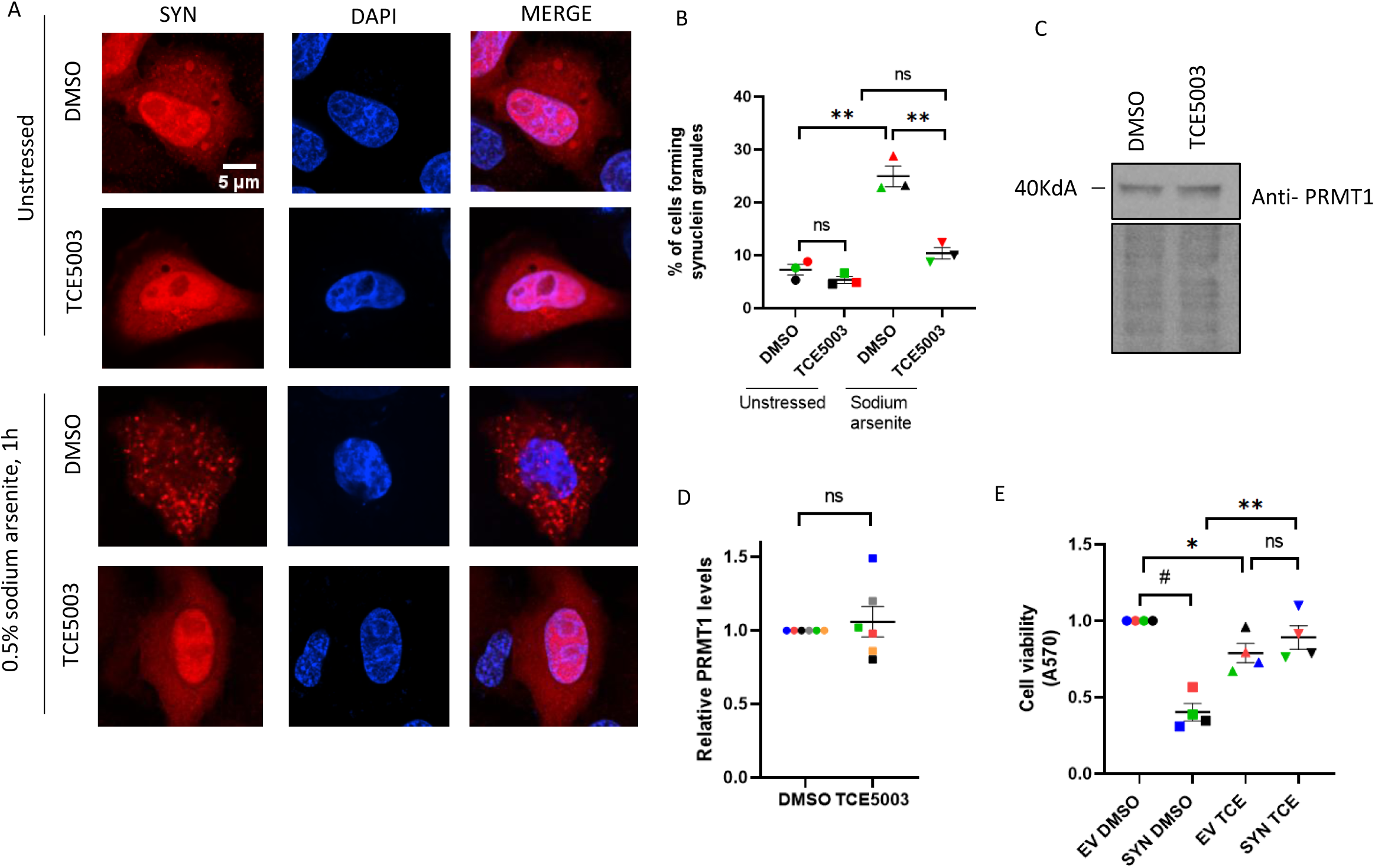
PRMT1 inhibitor (TCE5003) decreases Synuclein aggregates in HeLa cells. **A.** Microscopy for SYN upon addition of PRMT1 inhibitor-TCE5003 (5µM for 3h). **B.** Quantitation for data represented in A as % of cells forming SYN granules. **C.** western blotting analysis to check PRMT1 levels upon addition of PRMT1 inhibitor TCE5003. **D.** Quantitation for data represented in C. **E**. MTT assay for checking SYN toxicity upon addition of PRMT1 inhibitor. The significance of the data was calculated by paired t-test, and P-values are summarized as follows: ***P < 0.005; **P < 0.01; *P < 0.05.

**Figure 7.**
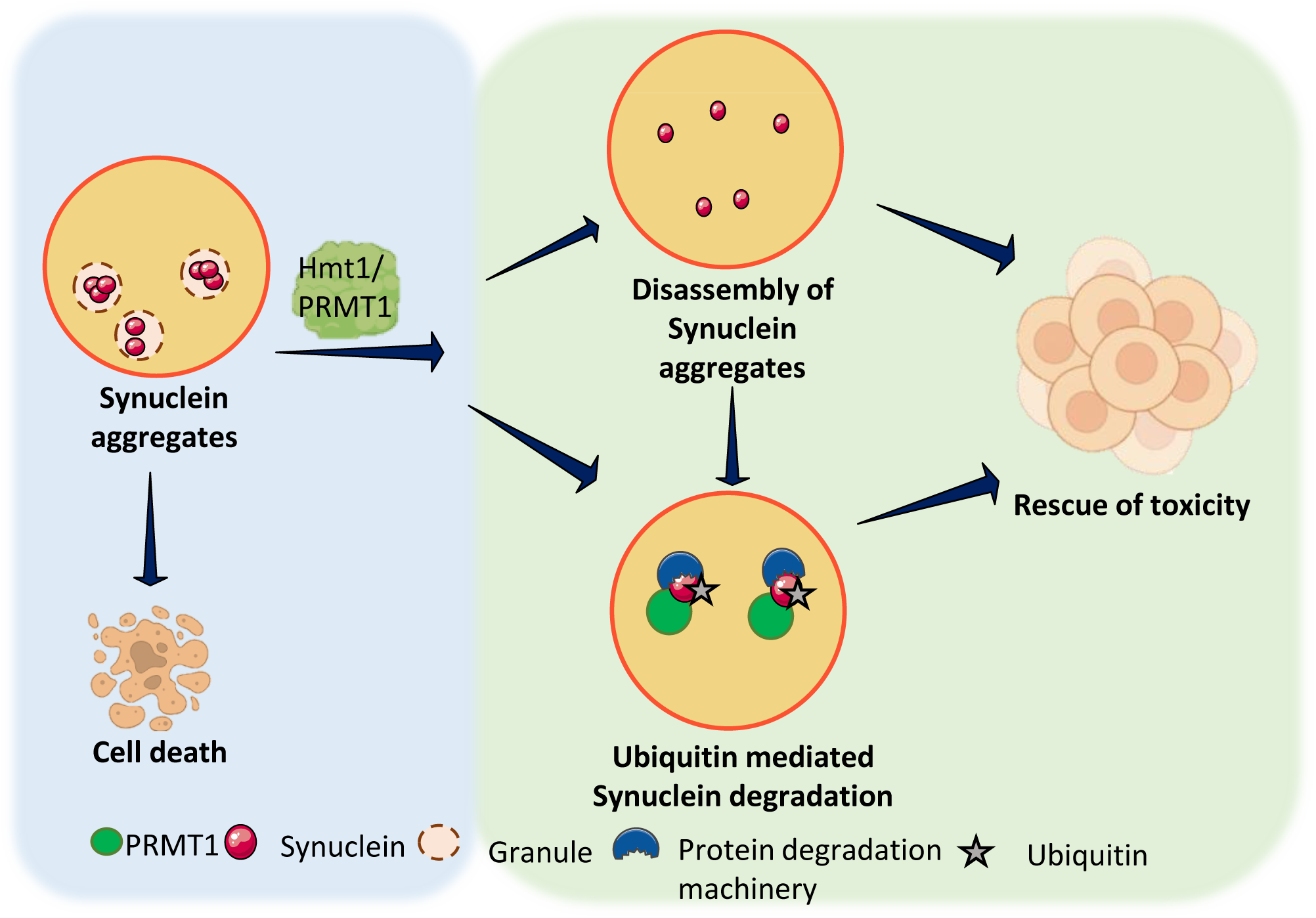
:PRMT1 acts as a disassembly factor for Synuclein. Hmt1/ PRMT1 can disassemble Synuclein aggregates and further maintain Synuclein protein levels via ubiquitination, which leads to a decrease in toxicity and rescue of cell growth

## Discussion

In this study, we describe a previously unrecognized role for the major arginine methyltransferase Hmt1 in yeast and its human homolog PRMT1 in regulating the aggregation and toxicity of α-synuclein. These enzymes have been studied for their roles in methylating arginine residues and in regulating gene expression. However, our work reveals that they also play an important part in helping cells cope with stress-induced protein aggregation. Several observations support this idea – (i) Hmt1/ PRMT1, a majorly nuclear protein, delocalises from nucleus to cytoplasm under oxidative stress, independent of its catalytic activity (Figure 1,4), (ii) Hmt1 attenuates α-synuclein toxicity by promoting its ubiquitination-dependent degradation and thereby regulating α-synuclein protein levels (Figures 2). (iii) α-synuclein aggregates increase in Δhmt1 and PRMT1 knockdown cells (Figure 3), (iv) PRMT1 suppresses α-synuclein granule formation in HeLa cells (Figure 4), (v) addition of purified Hmt1 and PRMT1 reduced α-synuclein in the granule-enriched fraction. (Figure 3&5), (vi) α-synuclein granule disassembly precedes the reduction in α-synuclein protein levels (Figure 3&5), (vii) PRMT1 inhibitor (TC-E-5003) decreases α-synuclein granules and associated toxicity in HeLa cells (Figure 6). These observations indicate that Hmt1/ PRMT1 is involved in disassembling α-synuclein granules (Figure 7).

Although stress-induced cytoplasmic relocalization of Hmt1 has not been previously reported, several studies have documented dynamic changes in PRMT1 localization under stress conditions. Studies have shown that PRMT1 partially co-localizes with FUS in cytoplasmic inclusions following arsenite-induced stress^18^. PRMT1 splice variant 2, which contains a nuclear export signal, exhibits high nucleocytoplasmic mobility, and inhibition of methyltransferase activity leads to its nuclear accumulation^19^. Notably, further studies demonstrated that a catalytically inactive mutant of PRMT1 splice variant 2 forms nuclear granules even under unstressed conditions^20^. In addition, PRMT1 accumulation has been observed in cytoplasmic bodies proximal to the nucleus, with γ- and UV-irradiation reducing the size of these structures^21^. It has also been reported that PRMT1 overexpression led to cytoplasmic PRMT1 granule formation, which colocalised with stress granules^22^. However, the functional relevance of PRMT1 localization to the cytoplasm remains unclear.

Our data show that both Hmt1 and PRMT1 relocalize from the nucleus to the cytoplasm in response to oxidative stress (Figures 1A and 4A), a condition known to promote protein damage and aggregation. Importantly, this relocalization occurs even when the enzymes are catalytically inactive (Figure 1E), suggesting that their stress-induced cytoplasmic localization is independent of methyltransferase activity. Together, these findings support a model in which Hmt1/PRMT1 act as stress-responsive factors whose physical presence in the cytoplasm—rather than their enzymatic activity—is central to the function we observe.

Oxidative stress is known to induce lipid peroxidation, protein oxidation, DNA and RNA damage in cells. Excessive generation of reactive oxygen species (ROS) and reactive nitrogen species (RNS) disrupts mitochondrial function and is strongly implicated in the pathogenesis of neurodegenerative disorders^23,14^. Parkinson’s disease is a prominent example, in which oxidative stress and mitochondrial dysfunction due to mutation in certain mitochondrial genes promote α-synuclein misfolding and the formation of toxic oligomeric species that drive neuroinflammation^24^. Under physiological conditions, α-synuclein plays an important role in lipid binding and regulates neurotransmitter release at synaptic junctions^25^.

*Saccharomyces cerevisiae* has emerged as a simple yet powerful model system for studying neurodegenerative disorders, including Parkinson’s disease^26,27^. Overexpression of α-synuclein in yeast is well known to induce cytoplasmic aggregation and cellular toxicity^28^. Consistent with previous reports, we observed robust α-synuclein–mediated toxicity and granule formation in yeast (Figure 2&3). Although multiple yeast-based genetic screens have identified modifiers of α-synuclein toxicity^29,13^, the potential role of these factors in actively disassembling α-synuclein granules has not been explored.

A recent study reported a genetic interaction between Hmt1 and α-synuclein, in which deletion of HMT1 enhanced α-synuclein–induced toxicity^13^. Based on this observation, we hypothesized that Hmt1 functions as a disassembly factor for α-synuclein aggregates. α-synuclein lacks arginine residues. Deletion of HMT1 increased cellular sensitivity to α-synuclein toxicity (Figure 2A) and led to a higher protein level of α-synuclein (Figure 2C). Complementation with plasmid borne Hmt1 rescued both the phenotypes (Figure 2B&C). Strikingly, a catalytically inactive Hmt1 mutant was even more effective than the wild-type protein in suppressing α-synuclein toxicity and protein levels (Figure 2B&C). We believe that as the catalytically dead mutant is free from performing its usual function, it might be more available for binding to Synuclein and thus shows a better rescue of toxicity. These results demonstrate that Hmt1 regulates α-synuclein aggregation through a mechanism independent of its methyltransferase activity, revealing a previously unappreciated non-catalytic role for Hmt1 in maintaining proteostasis.

Cellular levels of α-synuclein are tightly regulated by two major protein quality control pathways: lysosomal autophagy and the ubiquitin–proteasome system (UPS)^30,31,32^. Disruption of this proteostatic balance, such as during oxidative stress or upon mutations affecting mitochondrial function, is known to promote α-synuclein accumulation and aggregation, a central pathological feature of Parkinson’s disease^33,34^.

In this context, our observations reveal a previously unrecognized role for Hmt1 in regulating α-synuclein turnover. Deletion of *HMT1* resulted in a markedly reduced rate of α-synuclein degradation (Figure 2E), accompanied by diminished association of α-synuclein with the proteasomal machinery (Figure 2H) and a corresponding decrease in α-synuclein ubiquitination (Figure 2I). These defects point to a selective impairment of the UPS-mediated clearance of α-synuclein in the absence of Hmt1, further suggesting that Hmt1 makes the proteasomal machinery available for Synuclein, which leads to a decrease in Synuclein protein levels.

Overexpression of α-synuclein is known to promote the formation of cytoplasmic granules, a phenotype that is further exacerbated under oxidative stress conditions^35,36^. Interestingly, we observed a marked increase in α-synuclein granules formation in Δ*hmt1* cells (Figure 3A). Notably, these granules persisted even during the recovery phase following stress removal, in contrast to wild-type cells where granules were largely resolved (Figure 3A&C). This persistence suggests that Hmt1 contributes to the efficient clearance or disassembly of α-synuclein assemblies. Together, these observations support a role for Hmt1 in facilitating the dynamic remodeling of α-synuclein granules during stress recovery, thereby limiting their accumulation and associated cellular toxicity.

Our experiments provide mechanistic insight into how Hmt1 influences α-synuclein granule disassembly. In biochemical granule-enrichment assays, the addition of recombinant-purified Hmt1 led to a marked reduction in α-synuclein in the granule-enriched fraction, indicating a direct effect on α-synuclein granules. Notably, this activity was further enhanced by the catalytically inactive G68R mutant, reinforcing the idea that Hmt1’s role in this context is independent of its methyltransferase activity. These observations support a model in which Hmt1 directly engages α-synuclein granules and promotes their dissolution, thereby functioning as a disassembly factor (Figure 3F).

Analysis of overlap between known chaperone proteins in yeast^38^ and Hmt1 interacting proteins (from *Saccharomyces* genome database), indicated 3 common proteins (Supplementary Figure 2A). This suggests that Hmt1 could regulate the aggregation of aggregation prone proteins. Analysis of overlap between aggregation prone intrinsically disordered proteins in yeast^37^ (from DisProt) and Hmt1 interacting proteins (from *Saccharomyces* genome database), indicated 10 common proteins (Supplementary Figure 2B). This suggests that Hmt1 could act as a co-chaperone for recruiting chaperone proteins and thus regulate the misfolded proteins under oxidative stress (Supplementary Figure 2C). Molecular chaperones are well established as key regulators of α-synuclein aggregation and disassembly^39,40,41^. In line with this, our data indicate that Hmt1 functionally interfaces with the chaperone network. We find that Hmt1 reduces the partitioning of the Hsp40 co-chaperone, Sis1 into granule-enriched fractions. This observation suggests that Hmt1 may influence the chaperone landscape associated with α-synuclein assemblies. Hmt1 may function as a co-chaperone that facilitates or stabilizes the recruitment of canonical chaperones to α-synuclein granules, thereby promoting their disassembly. Such a role would be consistent with our broader findings that Hmt1 regulates α-synuclein aggregation independently of its methyltransferase activity, and supports a model in which Hmt1 acts as a scaffold or modulator to coordinate protein quality control machinery at sites of aggregation.

Importantly, we observed that the reduction in α-synuclein granules precedes a detectable decrease in total α-synuclein protein levels (Figure 5D). This temporal separation suggests that disassembly of α-synuclein granules is an early and primary event, occurring prior to protein clearance. We propose that Hmt1-mediated disassembly converts aggregated α-synuclein into more soluble or accessible species, which may subsequently become competent for downstream degradation pathways. Together, these findings place Hmt1 upstream of α-synuclein turnover, highlighting a previously unrecognized, non-enzymatic role for Hmt1 in regulating the aggregation of aggregation-prone proteins.

Motivated by these observations, we extended our analysis to mammalian cells. Consistent with previous reports, PRMT1 delocalized from the nucleus to the cytoplasm upon oxidative stress (Figure 4A). Several recent studies have highlighted a role for PRMT1 in regulating both physiological and pathological cytoplasmic assemblies. Studies have shown that PRMT1 mediated arginine methylation markedly suppresses TDP-43 aggregation^35,42^, while conditional expression of PRMT1 attenuates FUS-ΔC–induced stress granule (SG) formation^18^. Similarly, elevated levels of PRMT1 or PRMT5 decrease G3BP1-positive SGs^22^, whereas PRMT1 depletion has the opposite effect^18^. PRMT1 knockdown reduces the association of RAP55A with P-bodies^43^, and loss of PRMT1 enhances, while its overexpression suppresses, UBAP2L-positive SG formation^44^. In all these cases, the effects of PRMT1 on granule dynamics were attributed to its arginine methyltransferase activity.

In contrast, our findings reveal a previously unappreciated, methylation-independent function of PRMT1 in modulating cytoplasmic α-synuclein granules. We observed that PRMT1 overexpression significantly decreased α-synuclein granules (Supplementary Figure 5A), whereas PRMT1 knockdown led to their accumulation (Figures 4E). Moreover, PRMT1 physically associates with α-synuclein (Figure 5E) and limits its enrichment in granule-partitioned fractions (Figure 5A). Notably, this effect was further enhanced by a catalytically inactive PRMT1 mutant (Figure 5C), strongly supporting a non-enzymatic role for PRMT1 in the disassembly of α-synuclein granules.

Recent studies have examined the involvement of PRMT1 in neuronal survival and function in Parkinson’s disease (PD) models. PD is characterized by the selective loss of dopaminergic neurons in the substantia nigra and the accumulation of Lewy bodies composed primarily of aggregated α-synuclein^45,46^. In neurotoxic PD models induced by MPP⁺ or rotenone, both the expression and activity of PRMT1 are significantly increased, and elevated PRMT1 levels have been associated with enhanced apoptosis of dopaminergic neurons^47^. Mechanistically, PRMT1 has been shown to promote the nuclear translocation of apoptosis-inducing factor (AIF) by enhancing PARP1 overactivation, leading to energy depletion and DNA damage and ultimately triggering cell death in both in vivo and in vitro PD models^48,49,50^. In addition to PARP1–AIF signalling, PRMT1 may also influence dopaminergic neuron apoptosis through other pathways, including the regulation of apoptosis signal-regulating kinase 1 (ASK1)^51^. However, contrasting reports suggest a more complex role for PRMT1 in PD pathology. Elevated iron levels, which are commonly observed in the substantia nigra in PD, can induce oxidative stress and have been reported to reduce PRMT1 levels, concomitant with increased dopaminergic neuron death^52,53,54^. Together, these findings indicate that PRMT1 function in PD is context-dependent and may vary with the nature and extent of cellular stress.

Overexpression of α-synuclein is reported to be toxic to mammalian cells^55,16^. Consistent with the previous reports, we also observed increased toxicity upon α-synuclein overexpression (Figure 5G). Co-expression of PRMT1 with α-synuclein reduced α-synuclein induced cytotoxicity in mammalian cells (Figure 5G), and both wild-type and mutant PRMT1 rescued toxicity to a similar extent, further supporting a catalytic activity–independent role (Figure 5G). As seen earlier for yeast cells, we saw that granule disassembly begins before the overall protein levels decline in HeLa cells as well (Figure 5H, I&J).

PRMT1 inhibitors are currently being explored in clinical and preclinical settings for cancer therapy, primarily for their ability to suppress arginine methylation without necessarily altering PRMT1 protein abundance^56,57,58^. Leveraging this, we tested a PRMT1 inhibitor that selectively inhibits catalytic activity while preserving PRMT1 protein levels-TC-E-50003 ^57,17,59,60^. Treatment with this inhibitor resulted in reduction of α-synuclein granules (Figure 6A) and a concomitant improvement in cell survival (Figure 6E). These findings are consistent with our genetic and biochemical data and further support the notion that PRMT1 can modulate α-synuclein aggregation through a mechanism that does not rely on its methyltransferase activity. Instead, this suggests that the physical presence of PRMT1, rather than its enzymatic function, plays a key role in regulating α-synuclein granules disassembly and associated cytotoxicity.

Taken together, our results support a model in which Hmt1/PRMT1 act as stress-responsive factors that help disassemble α-synuclein granules and direct them toward degradation. This broadens our understanding of how cells maintain protein quality under stress.

Because abnormal aggregation of α-synuclein is a key feature of Parkinson’s disease, our findings could have implications for understanding disease mechanisms. Uncovering how Hmt1/PRMT1 contributes to aggregate clearance may point toward new strategies for reducing toxic protein assemblies in neurodegeneration.

## Materials and methods

### Yeast strains

All strains used in this study are listed in Table 1. Yeast strains were derived from BY4741 (wild type) and grown at 30 °C in either yeast extract/peptone medium or synthetic medium supplemented with appropriate amino acids and 2% glucose or galactose, as required. For secondary cultures, cells were diluted to an OD₆₀₀ of 0.1 and grown to mid-log phase (OD₆₀₀ = 0.5–0.6).

**Table 1:**
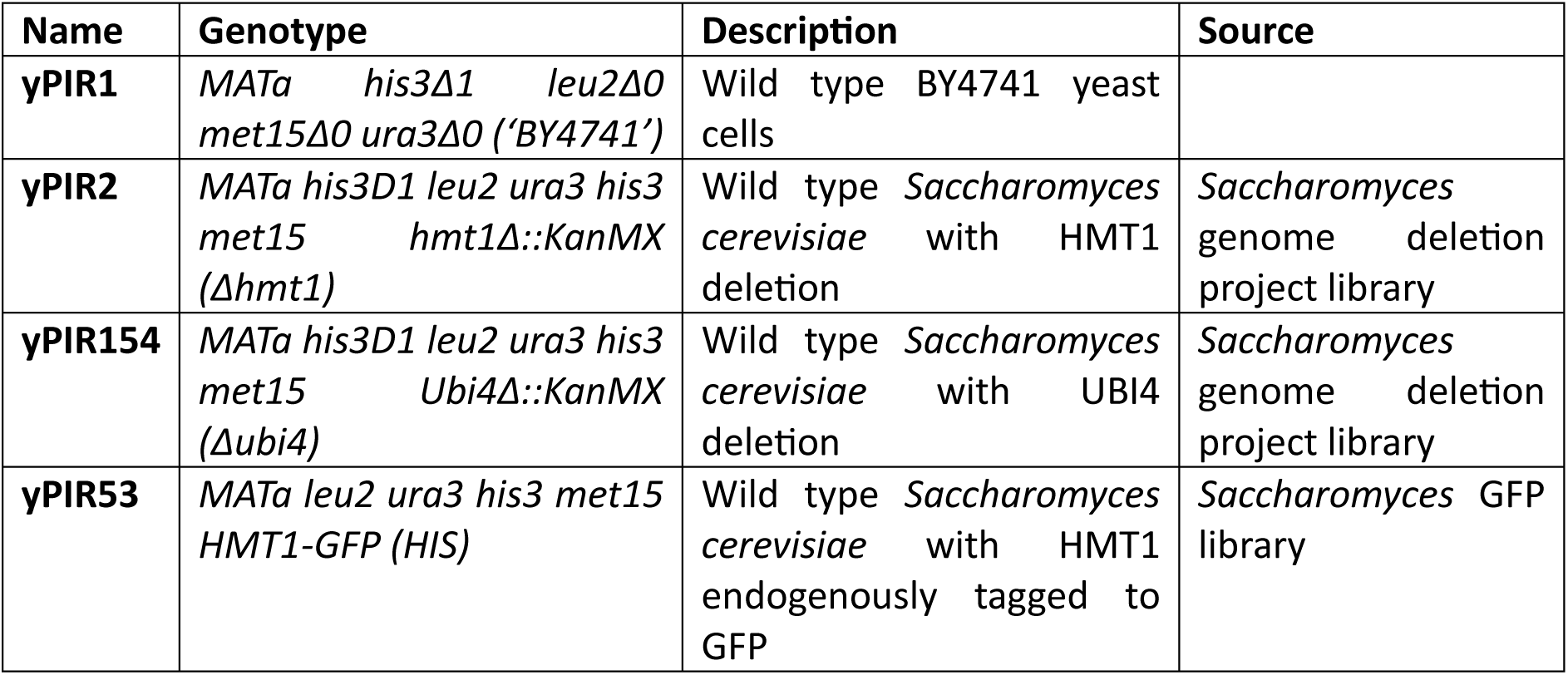
List of yeast strains used in this study.

For α-synuclein chase experiments, WT and Δhmt1 strains carrying the α-synuclein–GFP plasmid were grown in uracil-deficient medium containing 1% sucrose and 1% galactose to mid-log phase, then shifted to 2% glucose to repress expression. Cells were harvested after 1 and 2 h for analysis.

### Plasmid construction

The list of plasmids used in this study are listed in Table 2.

**Table 2:** List of plasmids used in this study.

| Name | Description | Source |
| --- | --- | --- |
| **pPIR353** | pRS315 plasmid expressing Hmt1- GFP under its own promoter, with *LEU2* and ampicillin resistance genes | This study |
| **pPIR354** | pRS315 plasmid expressing Hmt1 G68R- GFP under its own promoter, with *LEU2* and ampicillin resistance genes | This study |
| **pPIR76** | Plasmid expressing *hSYN-GFP* under a galactose inducible promoter, with *URA3* and ampicillin resistance genes | Addgene |
| **pPIR400** | Plasmid expressing *hSYN-mCherry* under a galactose inducible promoter, with *URA3* and ampicillin resistance genes | This study |
| **pPIR3** | pET28 *E. coli* expression vector for Hmt1 with N-terminal His Tag, Ampicillin resistance | This study |
| **pPIR425** | pET28 *E. coli* expression vector for Hmt1 G68R with N-terminal His Tag, Ampicillin resistance | This study |
| **pPIR365** | pcDNA vector expressing mScarlet under CMV promoter, hygromycin, and ampicillin resistance genes | Kind gift from Dr. Janin Lautenschlager |
| **pPIR390** | pcDNA vector expressing SYN-mScarlet under CMV promoter | Kind gift from Dr. Janin Lautenschlager |
| **pPIR264** | pGEX *E. coli* expression vector for PRMT1 with C-terminal GST Tag, Ampicillin resistance | Addgene |
| **pPIR52** | pRS416 plasmid expressing Edc3-mCherry under its own promoter, with *URA3* and ampicillin resistance genes | This study |
| **pPIR261** | pRS315 plasmid expressing Sbp1- mCherry under its own promoter, with *LEU2* and ampicillin resistance genes | This study |
| **pPIR96** | pYES plasmid expressing Scd6- mCherry under its own promoter, with *URA3* and ampicillin resistance genes | This study |
| **pPIR436** | pcDNA vector expressing PRMT1 WT-Myc under CMV promoter | Kind gift from Dr. Stephane Richard<sup>61</sup> |
| **pPIR437** | pcDNA vector expressing PRMT1 VLD-Myc under CMV promoter | Kind gift from Dr. Stephane Richard<sup>61</sup> |

### Yeast transformation

Cells were grown to an OD₆₀₀ of ∼0.5 in complete medium, harvested, washed once with water, and resuspended in 100 µl of 0.1 M LiAc. The suspension was aliquoted into 50 µl fractions and mixed with 240 µl of 50% (v/v) PEG, 36 µl of 1 M LiAc, 25 µl of salmon sperm DNA (100 mg/ml), and plasmid DNA. Samples were vortexed, incubated at 30 °C for 30 min, and heat-shocked at 42 °C for 15 min. Cells were then pelleted, resuspended in 100 µl water, and plated on glucose-containing minimal media. Plates were incubated at 30 °C for 2 days until colonies appeared.

### Stress treatment and recovery

Yeast strains were grown in SD–Leu medium containing glucose to an OD₆₀₀ of ∼0.6 and divided into equal aliquots. For sodium azide stress, cells were treated with 0.5% sodium azide for 30 min, with water as the vehicle control. For H₂O₂ stress, cells were treated with 4 mM H₂O₂ for 30 min, with water as the control. For nitrogen starvation, amino acid starvation, or glucose deprivation, cell pellets were resuspended in medium lacking nitrogen, amino acids, or glucose, respectively, and incubated for 30 min. For heat stress, cells were resuspended in pre-warmed medium and incubated at 42 °C for 30 min. For recovery experiments, stressed cultures were pelleted, washed three times with fresh medium containing the appropriate carbon source, and resuspended in fresh medium.

### Mammalian cell cultures

HeLa cells were maintained in Dulbecco’s Modified Eagle Medium (DMEM) supplemented with 10% fetal bovine serum (FBS) and 1× antibiotic–antimycotic solution (complete DMEM). Culture medium was replaced every 24 h, and cells were passaged upon reaching >90% confluency. Cultures were routinely tested for Mycoplasma contamination by a PCR-based assay on a monthly basis.

### Transfection and preparation of mammalian cell samples for microscopy analysis

Transfections were performed using Lipofectamine 2000 (Thermo Fisher Scientific) according to the manufacturer’s instructions. Cells were grown, transfected, and processed on coverslips. Twenty-four hours post-transfection, cells were fixed with 4% formaldehyde for 15–20 min, followed by three washes with 1× PBS.

For immunocytochemistry cells were permeabilized with 0.25% Triton X-100 for 25 min and blocked for 1.5–2 h in blocking buffer containing 1% BSA and 0.3% Triton X-100. Cells were then incubated with primary antibody (1:200 dilution in blocking buffer) overnight at 4 °C in a humidified chamber. The following day, cells were washed three times with PBST (PBS containing 1% Tween-20) and incubated with secondary antibody (1:500 dilution in blocking buffer) for 2 h at room temperature. After three additional PBST washes, nuclei were stained with DAPI. Coverslips were mounted using Fluoromount-G and stored at 4 °C until imaging.

### Microscopy analysis

For live-cell imaging of yeast, cells were harvested immediately after stress treatment by centrifugation at 14,000 rpm for 15 s, and 5 µl of the cell suspension was spotted onto No. 1 glass coverslips. Images were acquired using a DeltaVision RT microscope system running softWoRx 3.5.1 software (Applied Precision, LLC) with an Olympus 100× oil-immersion objective (NA 1.4). Images were captured as 512 × 512-pixel files using a CoolSnapHQ camera (Photometrics) with 1 × 1 binning and were deconvolved using standard softWoRx algorithms. At least 100 cells were quantified per experiment using ImageJ.

HeLa cells were imaged using the same DeltaVision RT system with an Olympus 60× objective. Image analysis was performed using Fiji–ImageJ. Granules were quantified using the *Find Maxima* tool after converting images to 8-bit format, with prominence set to 50, and the number of granules and cells was counted.

Total fluorescence intensity was quantified by measuring the corrected total cell fluorescence (CTCF) for defined regions of interest (ROI). Background fluorescence was calculated from three background regions per image, and the average background intensity was subtracted from the ROI signal using the following formula:

CTCF = (Area of ROI × Mean ROI intensity) – (Area of background × Mean background intensity)

For nuclear-to-cytoplasmic (N:C) ratio analysis, CTCF values for the nucleus and whole cell were calculated as described above. Cytoplasmic intensity was determined by subtracting nuclear CTCF from total cell CTCF, and the N:C ratio was then computed.

### Western blotting

SDS–polyacrylamide gels were run and proteins were transferred onto Immobilon-P membranes (Merck) using the Bio-Rad Trans-Blot Turbo system at 100 V for 90 min. Membranes were blocked in 5% skim milk prepared in TBST, incubated with primary antibodies overnight at 4 °C, followed by incubation with secondary antibodies for 1 h at room temperature. Blots were developed using Bio-Rad Western ECL reagents and imaged on a Bio-Rad ChemiDoc system. Antibodies used in this study are listed in Table 3.

**Table 3:**
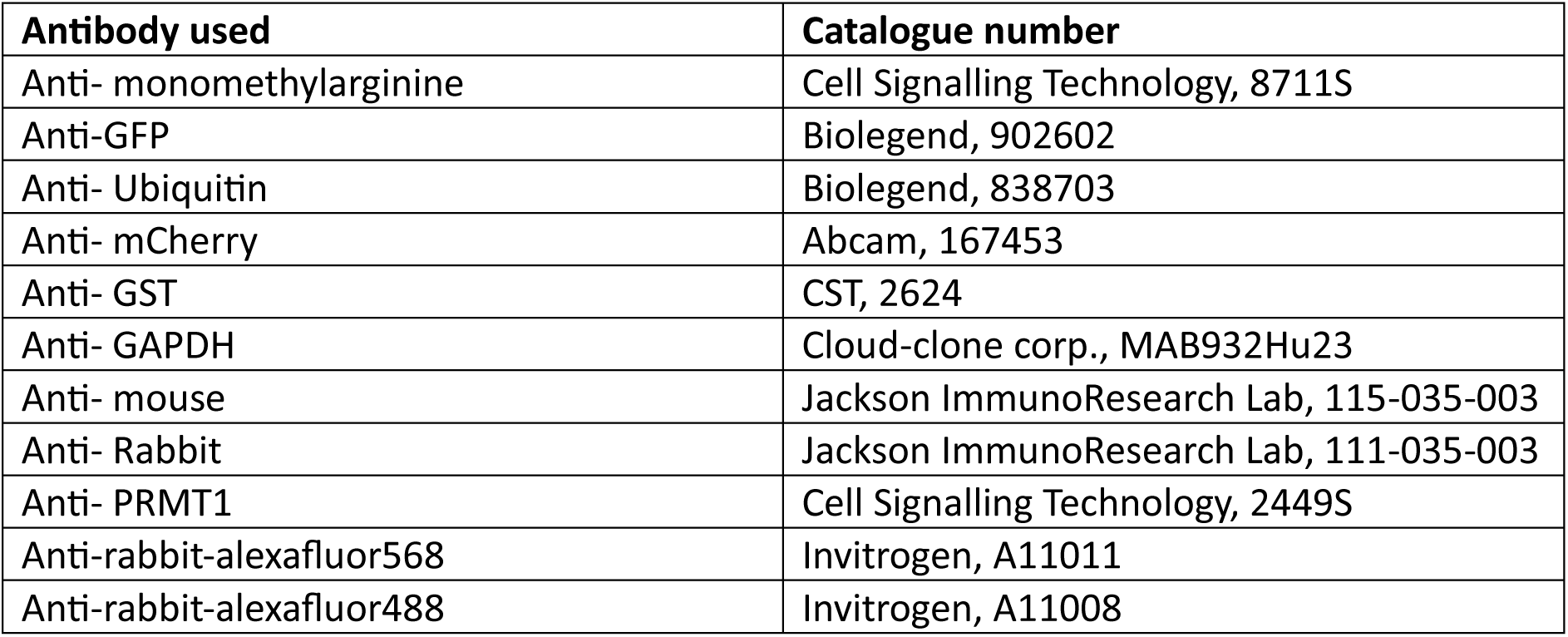
List of antibodies used in this study.

### Affinity pull-down for α-synuclein

For GFP-Trap pull-down assays, WT and Δhmt1 yeast strains expressing α-synuclein–GFP were grown in uracil-deficient medium containing 1% sucrose and 1% galactose to mid-log phase (OD₆₀₀ = 0.5–0.6). Cells from 100 ml cultures were harvested and lysed in 1 ml lysis buffer (10 mM Tris-Cl pH 7.5, 0.5 mM EDTA, 150 mM NaCl) by bead beating at 4 °C. Cell debris was removed by centrifugation at 5,500 rpm for 10 min at 4 °C.

For each pull-down, 1,000 µg of total protein was incubated in 1 ml buffer containing lysis buffer components, 1× protease inhibitor cocktail, and 5 µl GFP-Trap beads (Chromotek; cat. no. gtmak-20). After removing input samples, reactions were nutated at 4 °C for 2 h. Beads were washed three times (5 min each) with lysis buffer and resuspended in 50 µl dilution buffer. SDS sample buffer (10 µl) was added, and samples were boiled for 5 min. Approximately 6% input and 50% bead-bound fractions were analyzed by SDS–PAGE followed by western blotting using anti-ubiquitin antibodies. Total α-synuclein–GFP levels were assessed by re-probing the same blot with anti-GFP antibodies.

### Growth curve analysis

Overnight cultures of WT and Δhmt1 strains expressing α-synuclein–GFP were diluted to an OD₆₀₀ of 0.1 in SD–Ura medium containing 2% galactose. OD₆₀₀ readings were recorded every hour, and growth curves were plotted and analysed using GraphPad Prism 8.

### α-synuclein granule enrichment from yeast cells

WT and Δhmt1 strains expressing α-synuclein–GFP were grown in uracil-deficient medium containing 1% sucrose and 1% galactose to mid-log phase (OD₆₀₀ = 0.5–0.6). Cells from 100 ml cultures were harvested and lysed in 1 ml lysis buffer (10 mM Tris-Cl pH 7.5, 0.5 mM EDTA, 150 mM NaCl) by bead beating at 4 °C. Cell debris was removed by centrifugation at 2,000 × g for 2 min at 4 °C.

Lysates were divided into equal fractions, and ∼800 µg of protein was used for granule enrichment. Purified His–Hmt1 WT or G68R (6 µg), or an equivalent volume of buffer control, was added and incubated at 30 °C for 1 h. Samples were then centrifuged at 10,000 × g for 10 min at 4 °C to separate the supernatant (cytoplasmic fraction) and pellet (granules-enriched fraction), which were analysed by western blotting.

### α-synuclein granule enrichment from mammalian cells

HeLa cells transfected with α-synuclein–mScarlet were harvested 24 h post-transfection and lysed in RIPA buffer (50 mM Tris-HCl, 1% NP-40, 150 mM NaCl, 1 mM EDTA, 1 mM sodium orthovanadate, 1 mM sodium fluoride, 1× protease inhibitor cocktail, 1 mM PMSF, and RNase inhibitor) for 30 min at 4 °C. Cell debris was removed by centrifugation at 2,000 × g for 2 min at 4 °C.

Lysates were divided into equal fractions, and ∼800 µg of protein was used for granule enrichment. Purified PRMT1–GST (6 µg) or an equivalent volume of buffer control was added and incubated at 30 °C for 1 h. Samples were then centrifuged at 15,000 × g for 30 min at 4 °C to separate the supernatant (cytoplasmic fraction) and pellet (granule-enriched fraction), followed by western blot analysis.

### Expression and purification of recombinant proteins His-Hmt1 purification

His–Hmt1 plasmid was transformed into *E. coli* BL21 competent cells. Overnight cultures were diluted to an OD₆₀₀ of 0.1 in LB medium containing ampicillin and grown to OD₆₀₀ ∼0.5, followed by induction with 1 mM IPTG for 4 h at 37 °C. Cells were harvested by centrifugation at 4,200 rpm for 10 min at 4 °C, and pellets were stored at −80 °C until further use.

Cell pellets were resuspended in lysis buffer (50 mM NaH₂PO₄, pH 8.0; 300 mM NaCl; 10 mM imidazole; 1 mg/ml lysozyme; 1 µg/ml RNase A; 1 mM DTT; and 1× protease inhibitor cocktail) and incubated on ice for 30 min with intermittent mixing. Lysates were sonicated for nine cycles (10 s on/10 s off at 35% amplitude) and clarified by centrifugation at 15,000 rpm for 15 min at 4 °C.

The supernatant was incubated with Ni–NTA agarose beads (1:1 slurry, pre-equilibrated in lysis buffer) for 2 h at 4 °C with nutation. Beads were collected by centrifugation at 1,500 rpm for 1 min at 4 °C and washed three times with wash buffer (50 mM NaH₂PO₄, pH 8.0; 300 mM NaCl) containing increasing concentrations of imidazole (20 mM, 35 mM, and 50 mM), with each wash performed for 10 min at 4 °C.

Bound protein was eluted twice using elution buffer (50 mM NaH₂PO₄, pH 8.0; 300 mM NaCl; 500 mM imidazole) with nutation for 30 min at 4 °C. Eluted protein was dialyzed overnight at 4 °C against buffer containing 10 mM Tris (pH 7.0), 100 mM NaCl, 10% glycerol, and 1 mM DTT.

### PRMT1-GST purification

PRMT1–GST was expressed in *E. coli* BL21 cells grown in LB medium supplemented with ampicillin to an OD_600_ of 0.5, followed by induction with 1 mM IPTG for 4 h at 37 °C. Cells were harvested by centrifugation and stored at −80 °C until purification. For lysis, pellets were resuspended in lysis buffer (20 mM Tris-HCl, pH 8.0, 300 mM NaCl, 2 mM DTT) supplemented with lysozyme, RNase A, and protease inhibitor cocktail, and incubated on ice for 30 min. Cells were disrupted by sonication (9 cycles of 10 s on/off at 35% amplitude), and insoluble debris was removed by centrifugation at 15,000 rpm. The clarified supernatant was incubated with glutathione–Sepharose beads for 2 h at 4 °C. Beads were washed three times with high-salt wash buffer (500 mM NaCl), and bound protein was eluted twice with 20 mM reduced glutathione. The eluted protein was dialyzed overnight at 4 °C into storage buffer (10 mM Tris-HCl, pH 7.0, 100 mM NaCl, 10% glycerol, 1 mM DTT).

### MTT assay for mammalian cells

HeLa cells were seeded in 96-well plates and transfected with empty vector, α-synuclein–mScarlet, and PRMT1 wild-type or catalytically inactive mutant constructs. Twenty-four hours post-transfection, MTT reagent was added to each well to a final concentration of 0.5 mg/mL and incubated at 37 °C for 4 h. The MTT-containing medium was then removed, DMSO was added to solubilize the formazan crystals, and plates were incubated at 37 °C for 30 min. Absorbance was measured at 570 nm.

### Statistical analysis

All statistical analyses were performed using GraphPad Prism (version 8.0.2). Statistical significance was determined using unpaired or paired Student’s *t*-tests, or two-way ANOVA with multiple-comparison corrections, as indicated in the figure legends along with corresponding *p*-values. Error bars represent the standard error of the mean (SEM), and data points of the same colour within a graph denote values derived from the same experimental replicate.

## Acknowledgements

The authors thank Rajyaguru lab members for their inputs and suggestions during this work. We are grateful to Janin Lautenschlager for providing us with the Synuclein-mScarlet construct. We would also like to thank Stephane Richard for providing us with PRMT1 WT and PRMT1 VLD mutant constructs. We thank Priyanka Sanker for creating the Synuclein mCherry construct. PIR thanks the Department of Biotechnology [BT/PR51975/BMS/85/23/2024] for the funding. SD thanks CSIR for financial assistance.

---

**Figure S1:**
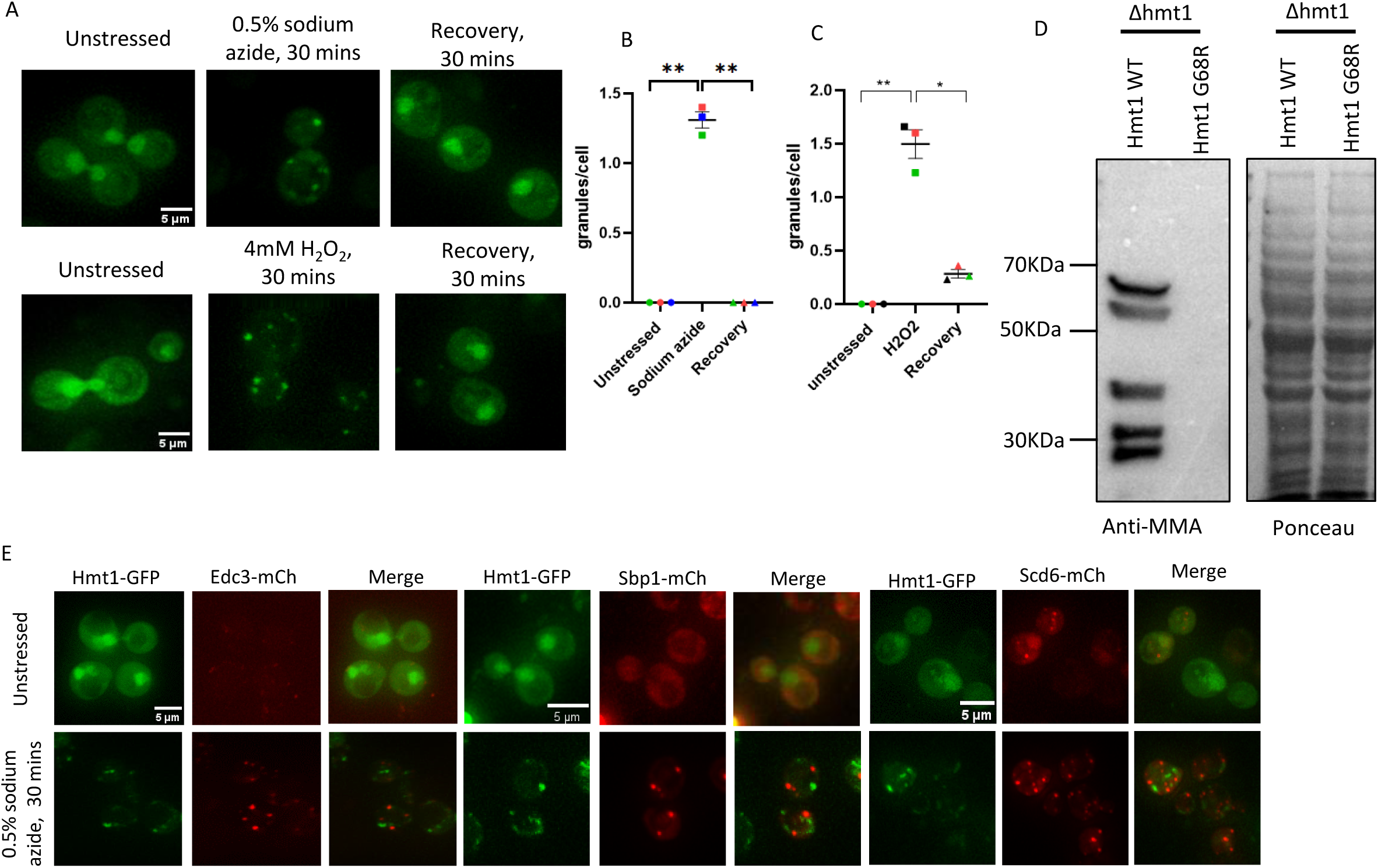
Hmt1 delocalises from nucleus to cytoplasm under oxidative stress. **A**. live cell microscopy for checking endogenous Hmt1-GFP localization under sodium azide and H_2_O_2_ stress stress. **B, C**. Quantitation for A as % of cells forming granules. **D**. Western analysis to check Hmt1 GFP levels upon treatment with 0.5% Sodium azide and 4mM H_2_O_2_ for 30 mins. **D.** Western blot for Hmt1 WT and Hmt1 G68R, probed with anti-mono methylarginine specific antibody showing G68R is catalytically dead. Ponceau serves as loading control. **E**. Live cell microscopy to check colocalization between -Hmt1 and P-body marker, Edc3, and between Hmt1 and its substrates-Sbp1 and Scd6, under unstressed and sodium azide stress. The significance of the data was calculated by paired t-test, and P-values are summarized as follows: ***P < 0.005; **P < 0.01; *P < 0.05.

**Figure S2:**
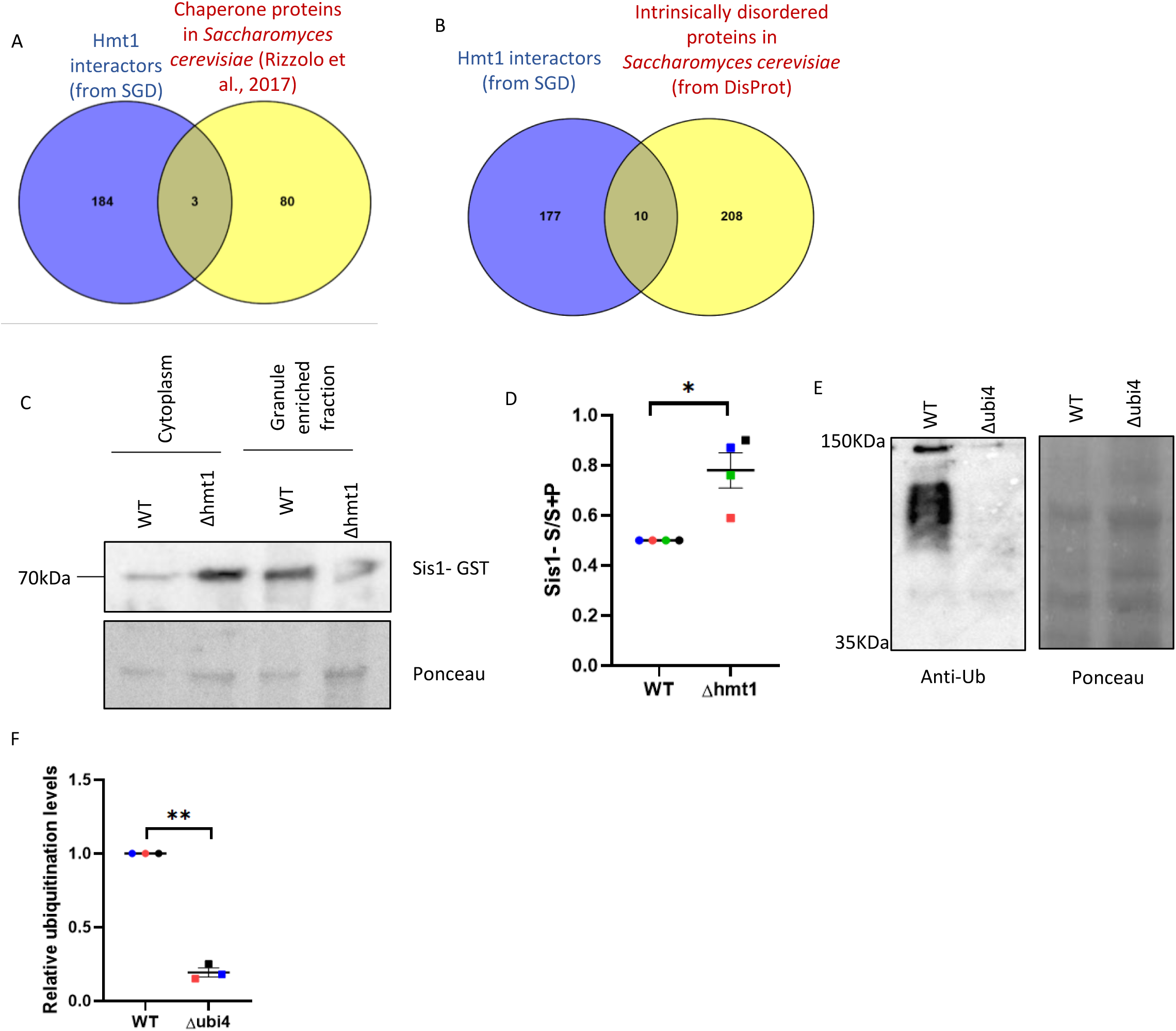
Hmt1 is involved in protein quality control pathway. **A.** Venn diagram to depict Hmt1 interacting proteins that could have chaperone activity. **B**. Venn diagram to depict Hmt1 interacting proteins that could have intrinsically disordered region. Venn diagrams were plotted using venny2.0 software. **C**. Granule enrichment for chaperone protein Sis1, showing decreased partitioning to granule enriched fraction upon deletion of Hmt1. **D**. Quantitation for data represented in C. **E**. Ubiquitination levels for WT and Δubi4, confirming that signal is specific to Ubiquitin. **F.** Quantitation. for E. The significance of the data was calculated by paired t-test, and P-values are summarized as follows: ***P < 0.005; **P < 0.01; *P < 0.05

**Figure S3:**
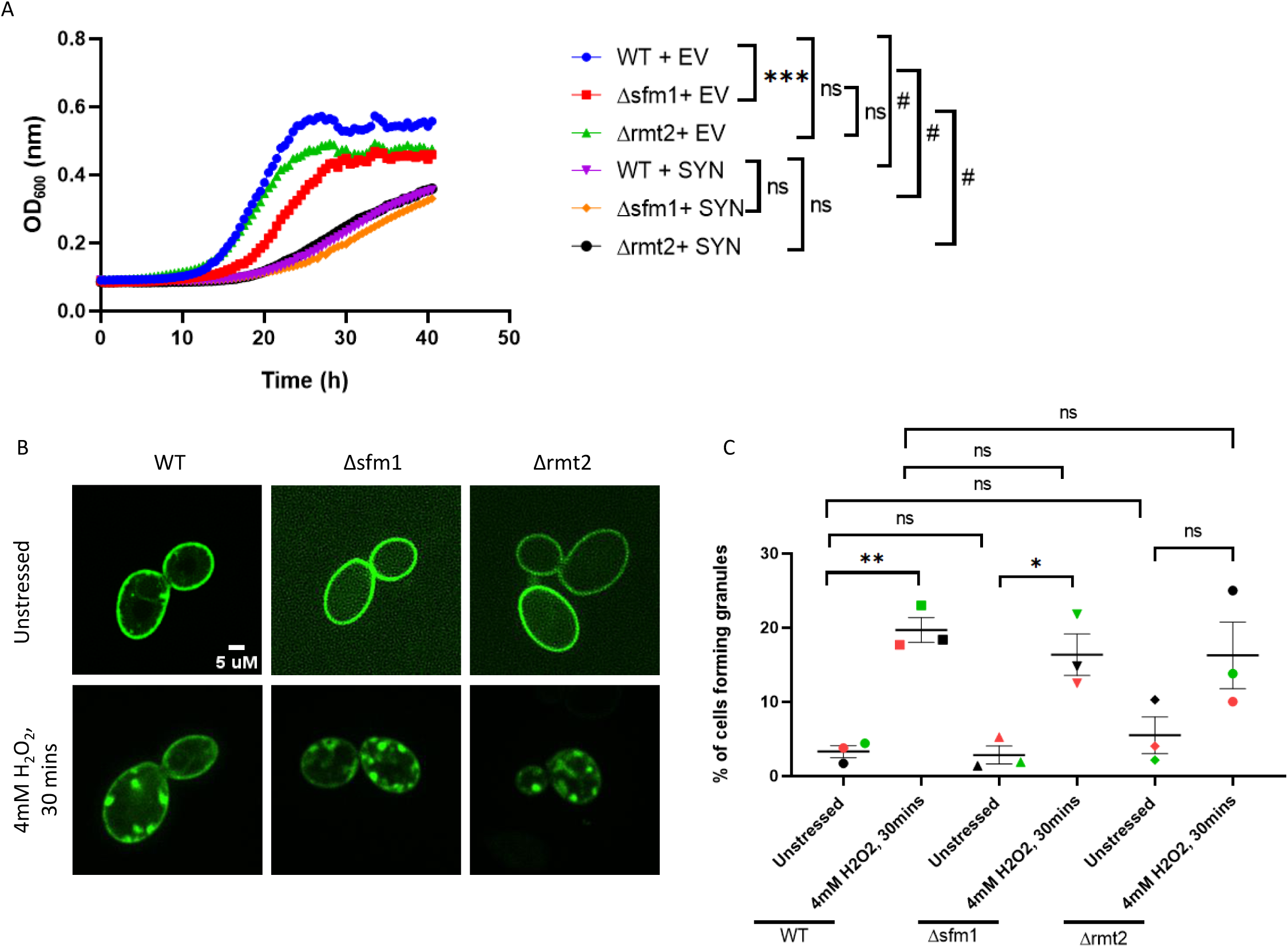
Deletion of other arginine methyl transferases, Sfm1 and Rmt2 does not affect Synuclein toxicity and localization to puncta. **A.** Growth curve analysis upon synuclein overexpression in WT, Δsfm1and Δrmt2 strains. Significance for growth curve data was calculated by two-way ANOVA. **B.** Live cell microscopy to check localization of Synuclein to puncta upon deletion of other arginine methyl transferases, Sfm1 and Rmt2, under unstressed and H_2_O_2_ stress. **C**. Quantitation for data represented in B as % of cells forming granules.

**Figure S4:**
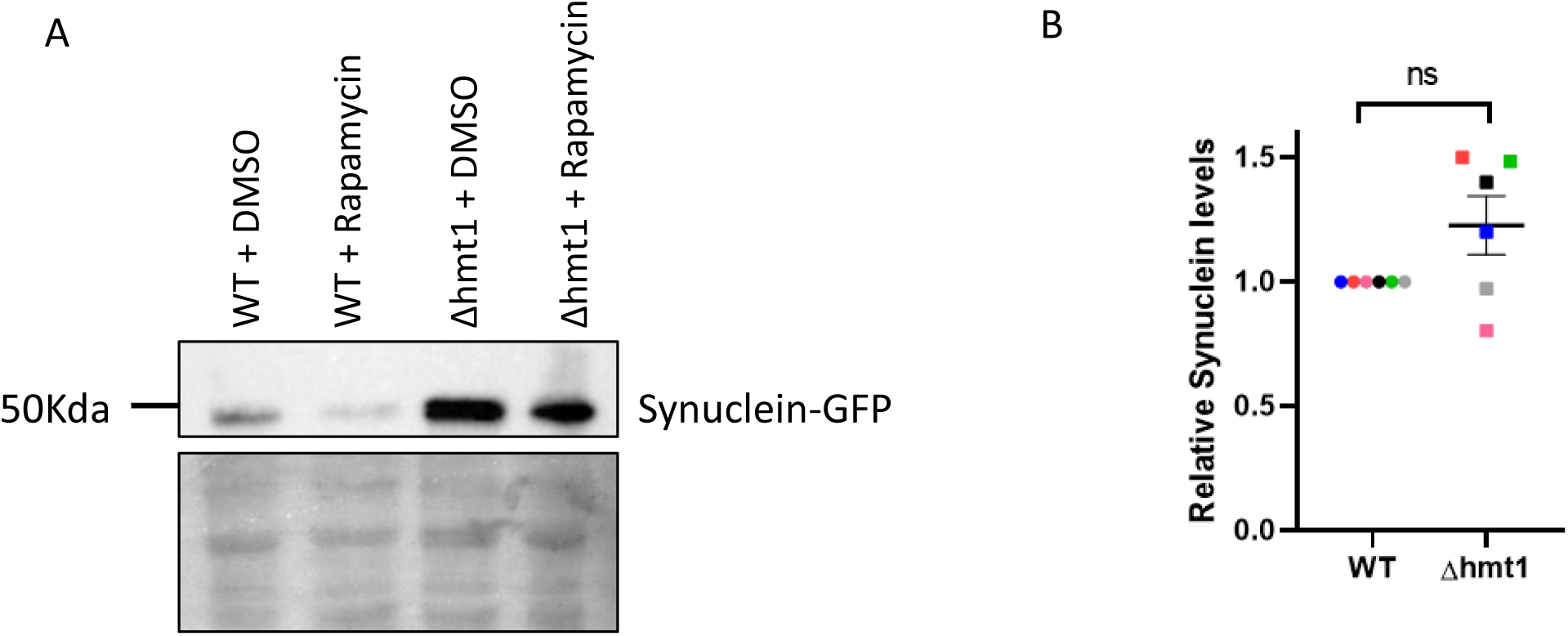
Hmt1 is not involved in autophagic degradation of Synuclein. **A**. Synuclein levels in WT and Δhmt1 upon treatment with Rapamycin and DMSO (Vehicle control). **B**. Quantitation for relative Synuclein levels upon Rapamycin treatment normalised w.r.t DMSO. The significance of the data was calculated by paired t-test, and P-values are summarized as follows: ***P < 0.005; **P < 0.01; *P < 0.05.

**Figure S5:**
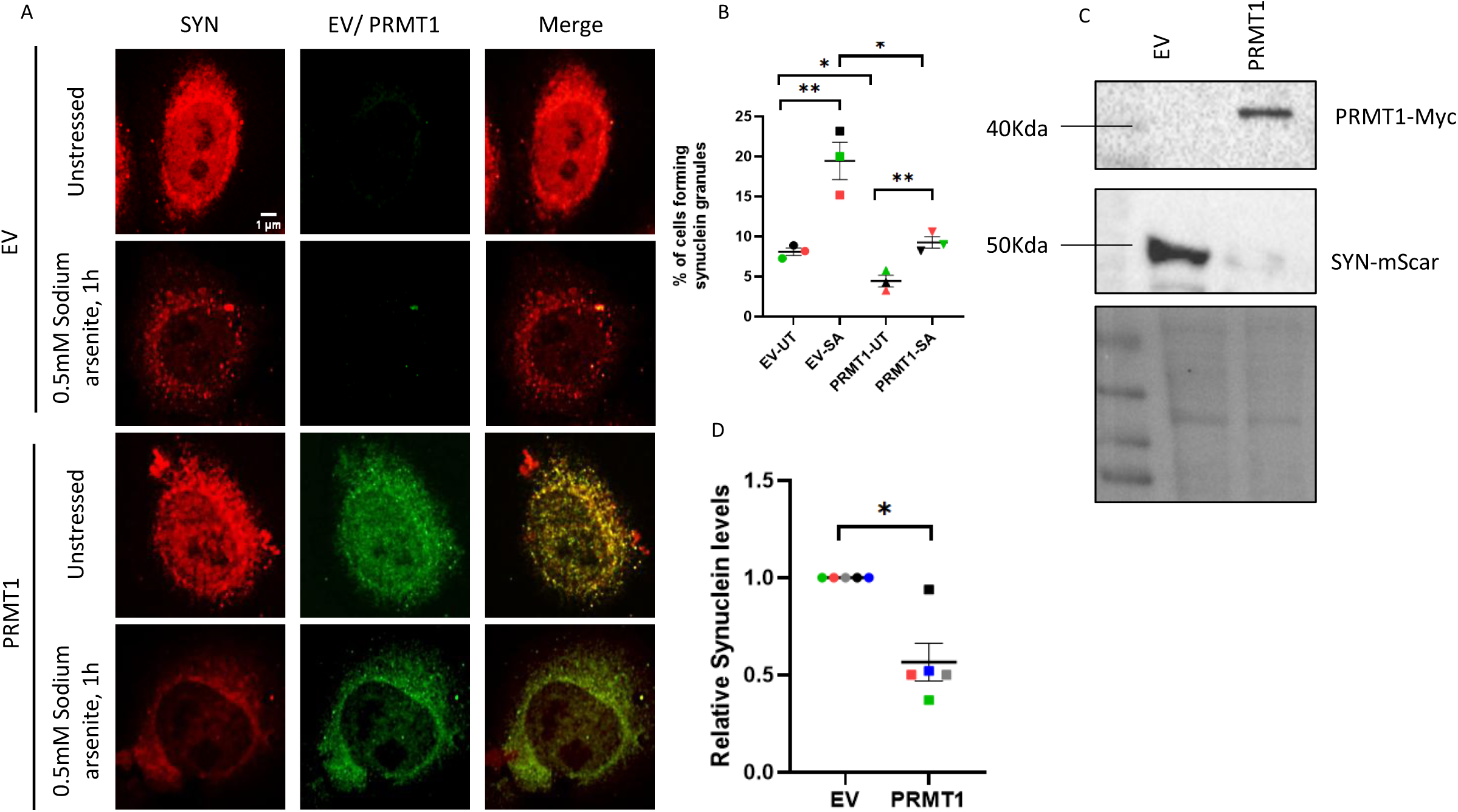
PRMT1 overexpression reduces Synuclein aggregates and protein levels in HeLa cells. **A.** Microscopy for SYN upon PRMT1 overexpression. **B.** Quantitation for A as % of cells forming granules. **C**. Western blotting analysis showing that overexpression of PRMT1 reduces SYN protein levels. **D.** Quantitation for data represented in C. The significance of the data was calculated by paired t-test, and P-values are summarized as follows: ***P < 0.005; **P < 0.01; *P < 0.05.

**Figure S6:**
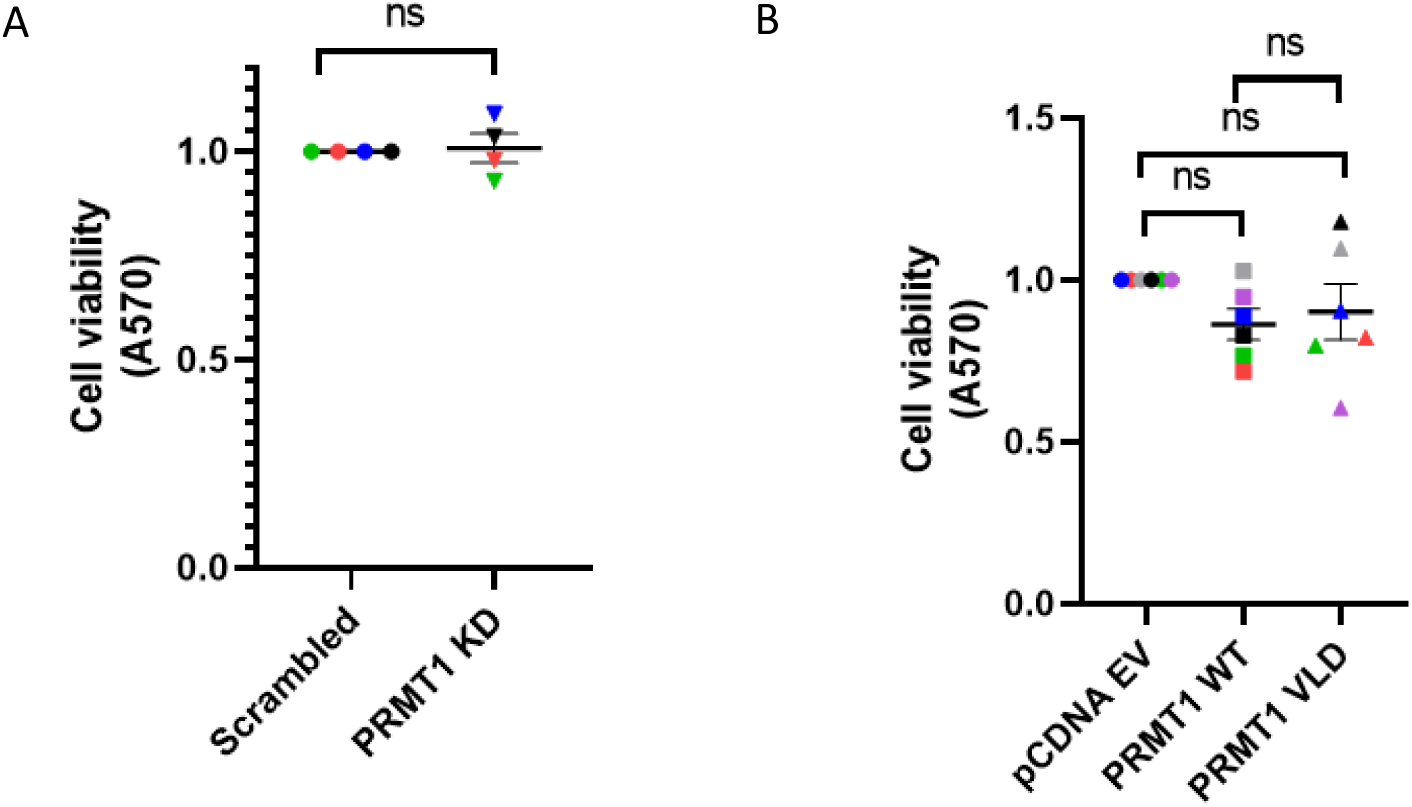
Knockdown and overexpression of PRMT1 itself does not affect cell viability. **A**.MTT assay upon PRMT1 knockdown. **B**. MTT assay for PRMT1 WT and catalytically dead mutant. The significance of the data was calculated by paired t-test, and P-values are summarized as follows: ***P < 0.005; **P < 0.01; *P < 0.05.

